# Predictive regulatory and metabolic network models for systems analysis of *Clostridioides difficile*

**DOI:** 10.1101/2020.09.14.297382

**Authors:** Mario L. Arrieta-Ortiz, Selva Rupa Christinal Immanuel, Serdar Turkarslan, Wei Ju Wu, Brintha P. Girinathan, Jay N. Worley, Nicholas DiBenedetto, Olga Soutourina, Johann Peltier, Bruno Dupuy, Lynn Bry, Nitin S. Baliga

**Affiliations:** Institute for Systems Biology, Seattle, WA, USA; Massachusetts Host-Microbiome Center, Dept. Pathology, Brigham & Women’s Hospital, Harvard Medical School, Boston, MA, USA; Université Paris-Saclay, CEA, CNRS, Institute for Integrative Biology of the Cell (I2BC), 91198, Gif-sur-Yvette, France; Laboratory of Pathogenesis of Bacterial Anaerobes, Institut Pasteur, Université de Paris, UMR-CNRS2001, France

## Abstract

Though *Clostridioides difficile* is among the most studied anaerobes, the interplay of metabolism and regulation that underlies its ability to colonize the human gut is unknown. We have compiled public resources into three models and a portal to support comprehensive systems analysis of *C. difficile*. First, by leveraging 151 transcriptomes from 11 studies we generated a regulatory model (EGRIN) that organizes 90% of *C. difficile* genes into 297 high quality conditional co-regulation modules. EGRIN predictions, validated with independent datasets, recapitulated and extended regulons of key transcription factors, implicating new genes for sporulation, carbohydrate transport and metabolism. Second, by advancing a metabolic model, we discovered that 15 amino acids, diverse carbohydrates, and 10 metabolic genes are essential for *C. difficile* growth within an intestinal environment. Finally, by integrating EGRIN with the metabolic model, we developed a PRIME model that revealed unprecedented insights into combinatorial control of essential processes for *in vivo* colonization of *C. difficile* and its interactions with commensals. We have developed an interactive web portal (http://networks.systemsbiology.net/cdiff-portal/) to disseminate all data, algorithms, and models to support collaborative systems analyses of *C. difficile*.

## INTRODUCTION

*Clostridioides difficile*, the etiology of pseudomembranous colitis, causes more than 500,000 infections, 30,000 deaths, and $5 billion per year in US healthcare costs (Monegro and Regunath, 2018). Infections arise through a variety of conditions that modulate the pathogen’s ability to colonize and expand in the gut. Antibiotic ablation of the commensal microbiota creates altered nutrient states in intestinal environments due to lack of competition for nutrients from host, dietary or microbial origin. The pathogen modifies its endogenous metabolism to respond to these altered states, which stimulates subsequent cellular programs that can promote enhanced colonization and growth. Stress and starvation conditions within *C. difficile* populations trigger responses that lead to sporulation, biofilm formation and release of mucosal damaging toxins (Aktories, 2011; Antunes et al., 2012; Saujet et al., 2011).

Symptomatic infection requires the production of toxins from the *C. difficile* pathogenicity locus (PaLoc), which includes the genes *tcdA, tcdB* and *tcdE* that respectively encode the A and B toxins and holin involved in toxin export (Govind and Dupuy, 2012). The PaLoc also contains *tcdR*, a sigma factor specific for the toxin gene promoters, and *tcdC*, a putative TcdR anti-sigma factor (Carter et al., 2011; Cartman et al., 2012; Mani and Dupuy, 2001; Matamouros et al., 2007). *C. difficile* elaborates toxin under starvation conditions to extract nutrients from the host and promote spore shedding (Edwards et al., 2016; Martin-Verstraete et al., 2016; Walter et al., 2014). Regulation of PaLoc expression occurs via a complex network of transcription factors (TFs) and small molecule inputs, of which direct primary regulators have been well described, but more complex and combinatorial effects remain unclear (Martin-Verstraete et al., 2016). Toxin production further triggers rapid and profound host immune responses, including release of reactive oxygen species which substantially alters the redox state of the gut environment, and other innate immune responses that can induce *C. difficile* stress responses to cell wall, oxidative, and other damaging stimuli (Bradshaw et al., 2017; Kint et al., 2017; Neumann-Schaal et al., 2018; Woods et al., 2016). As per all microbes, *C. difficile* adapts to complex, dynamic environments through changes in metabolism coordinated by a gene regulatory network (Brooks et al., 2011; Elena and Lenski, 2003). However, the mechanisms by which the gene regulatory network and metabolic pathways integrate to modulate *C. difficile* pathogenesis remain ill-defined (McDonald et al., 2018; Vemuri et al., 2017).

The *C. difficile* 630 (CD630) genome encodes 4,018 genes, with ∼309 candidate TFs (including sigma factors), 1,030 metabolic genes, and 1,330 genes (>30%) with unknown function (Monot et al., 2011; Riedel et al., 2015). The clinical ATCC43255 strain of *C. difficile,* used to capitulate symptomatic infections in mouse models, encodes 4,117 genes and ∼327 putative TFs, of which ∼97% are orthologous to genes encoded in the CD630 strain (Girinathan et al., 2021). To address questions regarding the broader systems-level interplay among genes in colonization and infection, we used computational modeling and network inference algorithms to construct an Environment and Gene Regulatory Influence Network (EGRIN) model for *C. difficile.* This model leverages a compendium of 151 published transcriptomes that surveyed responses of CD630 in diverse contexts. The EGRIN model consists of modules of putatively co-regulated genes identified based on their co-expression over subsets of conditions, enrichment of functional associations, chromosomal proximity, and presence of cis-acting gene regulatory elements (GREs) within their promoter regions indicating regulation by the same TFs. Further, using regression analysis, EGRIN also captures the combinatorial regulation of genes within each module as a function of the weighted influences of TFs. The model supports a systems-level understanding of the infective capacity of this obligate anaerobe under different *in vitro* and *in vivo* conditions.

In addition to EGRIN, we have advanced a metabolic network model of *C. difficile* to understand how conditional regulation manifests physiologically, by adding reactions and associated genes supporting the exchange of nutrients required for growth in intestinal environments. Integration of transcriptional and metabolic networks into a **P**henotype of **R**egulatory influences **I**ntegrated with **M**etabolism and **E**nvironment (**PRIME**) model supports prediction of conditional fitness contribution of every TF and metabolic gene of *C. difficile* (Immanuel et al., 2021). Analyses uncover TFs driving essential adaptive responses in *in vivo* conditions. This analytic framework provides a new systems-level view of the transcriptional and metabolic networks that coordinate *C. difficile’s* colonization, growth, expression of toxin, and adaptions to changing environments with host infection. Our models identified multiple TFs that coordinate critical aspects within each of these components, including contributions from PrdR, which regulates the Stickland proline and glycine reductase systems and other energy-generating pathways, and Rex a regulator modulating energy balance in *C. difficile* (Bouillaut et al., 2013, 2019). The PRIME model successfully identified enhanced epistasis between *ccpA* and *codY* in the presence of a protective gut commensal species. These findings refine the context and roles of these and other regulators in *C. difficile* virulence, and provide specific targets of vulnerability for model-informed interventions against this pathogen. The compiled datasets, algorithms, and models can be explored interactively through a community-wide web resource at http://networks.systemsbiology.net/cdiff-portal/.

## RESULTS

### Reconstruction of the environment and gene regulatory influence network (EGRIN) model for *C. difficile* 630

To investigate *C. difficile’*s transcriptionally-driven adaptive strategies we compiled 151 publicly available transcriptomes from 11 independent studies on CD630 (**Table S1**; *methods*). This compendium captures diverse transcriptional responses of *C. difficile* to commensals, *in vitro* and *in vivo* responses to different nutrient conditions, and consequences of targeted TF gene deletions. The transcriptome compendium together with functional associations information was analyzed with a suite of network inference tools (i.e., cMonkey2 and the Inferelator) to infer an EGRIN model for *C. difficile* (**Fig. 1A**; *methods*)(Arrieta-Ortiz et al., 2015; Reiss et al., 2015). The resulting EGRIN model organized 3,995 of 4,018 CD630 genes into 406 gene modules (using cMonkey2), and inferred module regulation (using the Inferelator) by 138 of 309 genomically identified TFs that putatively act through GREs discovered within gene and operon promoters. Among the Inferelator implicated regulatory networks, 255 modules were controlled by more than one TF, and 120 were regulated by more than two TFs (**Fig. S1**). The TF module assignments support subsequent hypothesis-driven experimental analyses, including the design of ChIP-seq and TF-deletion experiments to validate the regulatory network architecture under physiologically relevant environmental contexts.

**Figure 1.**
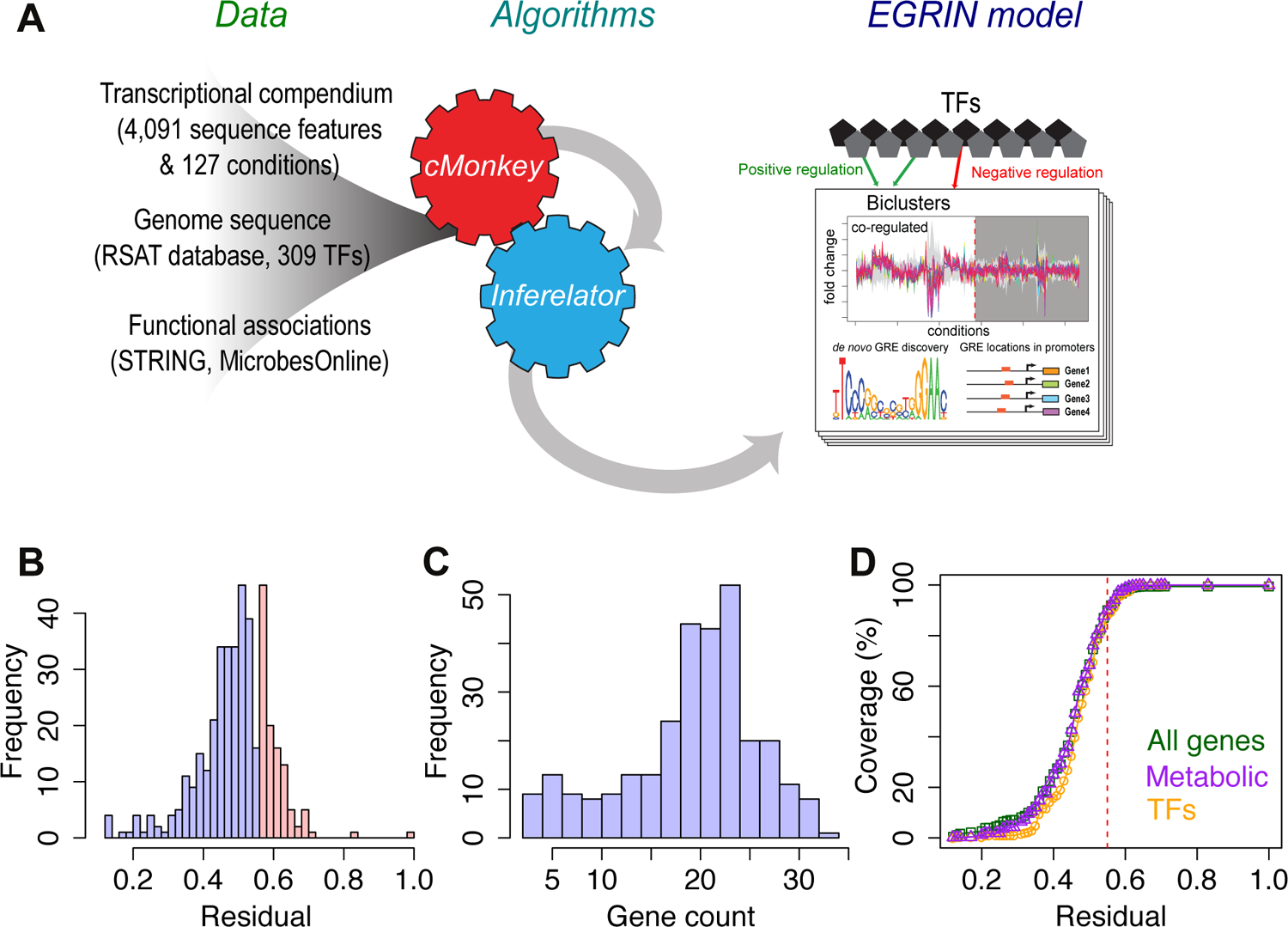
Inference pipeline and general properties of the resulting Environment and Gene Regulatory Influence Network (EGRIN) model of *C. difficile*. (A) Framework used to build the EGRIN model. (B) Distribution of residual values for the 406 detected co-regulated gene modules. 297 gene modules had residual values equal or lower than 0.55 (shown in purple) and were labelled as high quality. (C) Distribution of gene count for the high quality gene modules. (D) Coverage of all genes (4,018), the subset of metabolic genes (1,030) and TFs (309) by EGRIN modules for different residual thresholds. The red dashed line indicates the 0.55 residual cutoff.

**Table 1.** Experimental validation of PRIME-predicted synergistic epistasis between CcpA and CodY.

| TF<br>KO Strain | Mono-colonized |  |  |  | PBI co-colonized |  |  |  |
| --- | --- | --- | --- | --- | --- | --- | --- | --- |
|  | PRIME prediction |  | Experiment |  | PRIME prediction |  | Experiment |  |
|  | Relative growth (Log10)<sup>a</sup> | Essentiality<sup>b</sup> | Relative growth (Log10)<sup>a,c</sup> | Essentiality<sup>b</sup> | Relative growth (Log10)<sup>a</sup> | Essentiality<sup>b</sup> | Relative growth (Log10)<sup>a,c</sup> | Essentiality<sup>b</sup> |
|  |  |  | 24 h |  |  |  | 24 h |  |
| $\Delta ccpA$ | -0.716 | Essential | 0.609* | Non-essential | -0.144 | Non-essential | 0.315<sup>#</sup> | Non-essential |
| $\Delta codY$ | -0.482 | Essential | -1.181** | Essential | -0.176 | Non-essential | 0.527<sup>#</sup> | Non-essential |
| $\Delta ccpA \Delta codY$ | -1.198 | Essential | -1.299*** | Essential | -0.885 | Essential | -1.708*** | Essential |

<sup>a</sup>Relative growth with respect to wild type genotype
<sup>b</sup>Gene deletions that reduce *C. difficile* growth by more than 65% (i.e., log10(relative growth) < -0.456) were labelled as essential
<sup>c</sup>T-test P-values $\geq 0.05$ are indicated with the '#' symbol and were considered not significant.
The '\*', '\*\*' and '\*\*\*' symbols indicate P-values < 0.05, < 0.01 and < 0.001, respectively.

The quality of modules within the EGRIN model was evaluated using residual scores, which reflect the coherence of gene co-expression patterns. The lower the residual score, the higher the quality of the module. We determined using an empirical approach that a residual cutoff of 0.55 identified a functionally meaningful set of 297 high quality modules (73% of the total 406 modules) based on the relative enrichment of related functions within modules that passed filtering (**Fig. 1B**). This empirically determined threshold was similar to the threshold used to identify high quality EGRIN gene modules for *Mycobacterium tuberculosis* (Peterson et al., 2014). Altogether, the 297 high quality modules captured transcriptional regulation of 3,617 genes (90%) in CD630, with average membership of 20 genes per module (**Fig. 1C-D**). These metrics were consistent with models developed for other organisms (Brooks et al., 2014; Peterson et al., 2014), a remarkable finding given that the transcriptional dataset used to construct the *C. difficile* model was less than 10% the size of ones used to construct models for other species.

### Validation of the modular architecture and regulatory mechanisms uncovered by the *C. difficile* EGRIN model

We tested the accuracy of the EGRIN model to reconstruct previously characterized regulons and recapitulate key aspects of *C. difficile* biology. To do so, we performed gene enrichment analysis within modules using an updated annotation of *C. difficile* genome (Girinathan et al., 2021). This analysis identified 93 of 297 modules (31%) with significant enrichment of genes with related functions in 45 pathways (hypergeometric test adjusted p-value ≤ 0.05). Among these pathways, 14 were over-represented in three or more modules (**Fig. 2A**), demonstrating the capacity of the model to discover conditional partitioning of cellular processes. We also investigated whether the EGRIN model had identified known regulatory interactions between TFs and their target genes. We compiled from literature the regulons (i.e., target genes) of 13 previously characterized TFs in *C. difficile*, representing a network of 1,349 TF-gene interactions (**Table S2**). Notably, a total of 57 modules (19% of all high-quality modules) were significantly enriched with nine of these TF regulons (**Fig. 2B**). The EGRIN model recapitulated 514 of the 1,208 (42.5%) previously characterized interactions. This value is consistent with the recall rate of the EGRIN model for *M. tuberculosis* (41%-49%) (Peterson et al., 2014). The poor recall of the remaining four regulons (141 regulatory interactions) could be due to underrepresentation of gene expression data from relevant conditions in which these regulons are conditionally active. This analysis also uncovered combinatorial regulation of genes across 19 modules (i.e., enriched with more than one TF regulon). Consistent with the known hierarchical scheme for regulation of sporulation (Saujet et al., 2013), expression of 161 genes across at least eight modules were putatively influenced by Spo0A in combination with one or more alternative sigma factors implicated in sporulation (e.g., SigE). EGRIN also predicted CcpA contributions in six additional modules in combination with CodY, PrdR and SigL, illustrating the complexity of modular transcriptional regulation in *C. difficile*. The EGRIN modules also detected co-regulation of genes within and across functionally related operons. For example, module #152, which is enriched with the SigD regulon, contains 16 genes that are part of four operons including the flagellar operon *flgG1G*-*fliMN*-CD630_02720-*htpG,* in addition to *pyrBKDE*, CD630_30270-CD630_30280-*malY*-CD630_30300, and CD630_32430-*prdA* (**Fig. S2A**).

**Figure 2.**
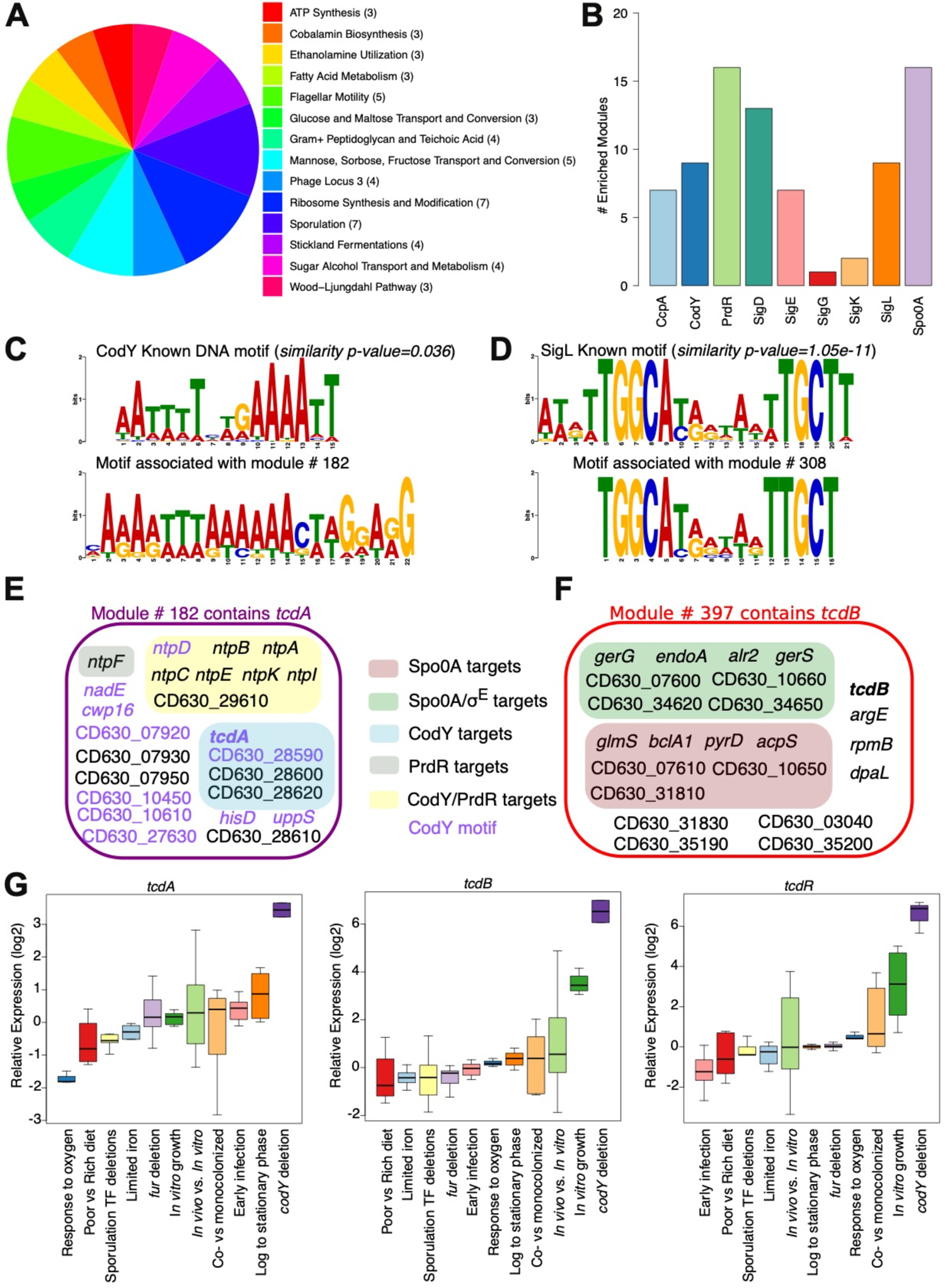
The Environment and Gene Regulatory Influence Network model of *C. difficile* recapitulates known biology of the pathogen. (A) Co-regulated gene modules are enriched with functional terms derived from expert curated annotation of the *C. difficile* genome (Girinathan et al., 2021). The pie chart shows terms over-represented in three or more modules. Number of modules associated with each functional term is shown in parenthesis. (B) Enriched gene modules among nine (out of 13) manually-defined and experimentally supported TF regulons (compiled from publicly available data in Table S2). (C) EGRIN identified the known DNA binding motif of CodY (Dineen et al., 2010). (D) EGRIN also identified the known DNA binding motif of SigL (Soutourina et al., 2020). Motif comparisons were performed using Tomtom (Bailey et al., 2015). (E) The EGRIN model recapitulated the previously reported influence of CodY on *tcdA* expression. The module #182 contains *tcdA*, it is enriched with members of the CodY regulon and contains a GRE (shown in panel C) similar to the experimentally determined CodY motif. (F) The EGRIN model also captured the interaction between toxin expression and sporulation via module #397 that contains *tcdB* and is enriched with genes regulated by sporulation-related transcriptional regulators. (G) Expression profiles of *tcdAB* and *tcdR* (positive regulator of the pathogenicity loci). Highest expression of the toxin genes and their activator was observed in the *codY* single deletion condition (dark purple box).

The clustering of genes in the EGRIN model is constrained by the *de novo* discovery of conserved GRE(s) within their promoters in order to cluster genes that are co-regulated, and not just co-expressed. The GREs represent putative binding sites for TFs that are often independently implicated by the Inferelator and protein-DNA interaction maps as regulators of genes within the same module (Bonneau et al., 2007; Brooks et al., 2014; Peterson et al., 2014). For instance, the regression-based analysis by the Inferelator assigned 138 TFs as putative regulators of the 297 gene modules, hypothesizing putative TF-GRE associations. While the paucity of characterized binding sites in *C. difficile* limited the degree to which these TF-GRE mappings could be validated, the predicted context-specific TF regulation of genes within modules can serve as a guide for performing ChIP-seq experiments in relevant conditions to iteratively improve the mechanistic accuracy of the EGRIN model. Notably, we determined that the GREs within promoters of genes in modules #182 and #308 recapitulated the known consensus binding sequences for CodY and SigL, which are predicted regulators of those modules, and are among the very few TFs in *C. difficile* for which binding sites have been characterized (**Fig. 2C-D** and **Fig. S2B-C**)(Dineen et al., 2007; Soutourina et al., 2020).

### *C. difficile* EGRIN model uncovers regulatory networks for the Pathogenicity Locus

We evaluated capacity for the EGRIN model to recall known mechanisms of PaLoc regulation, and to provide new information regarding complex regulatory and small molecule effects on toxin production. The EGRIN model captured certain previously described effects of CodY on toxin gene expression (**Fig. 2E**), as shown in module #182, which is enriched with members of the CodY regulon including *tcdA*. In agreement with EGRIN-predicted CodY regulation of PaLoc genes in module #182, genes encoding the toxin *tcdA* and its regulator *tcdR* were significantly overexpressed upon deletion of *codY* (**Fig. 2G**). Interestingly, *tcdB* (which is not part of module #182) was also significantly upregulated in the *codY* deletion strain, suggesting that this effect might be an indirect consequence of disrupted CodY regulation of *tcdR* (**Fig. 2G**). It has been assumed that CodY acts on PaLoc gene expression primarily through its repression of *tcdR.* Putative lower affinity binding sites have been suggested in the toxin gene promoter regions (Dineen et al., 2007). The presence of the CodY motif (**Fig. 2C**) in most members of module #182, including *tcdA* (purple font in **Fig. 2E**) suggests direct influence of CodY on *tcdA* gene expression. The EGRIN model also identified previously reported connections between sporulation and toxin production (Underwood et al., 2009). The *tcdB* toxin gene was assigned to module #397, which was significantly enriched with genes controlled by Spo0A, the master regulator of sporulation (**Fig. 2F**). Additional members of the PaLoc were assigned to other modules, supporting the presence of multiple condition-dependent promoters within the PaLoc (**Table S3**).

### Assignment of putative functions to genes in EGRIN modules

Approximately 33% of gene features in the CD630 genome have unknown functions. Thus, the *C. difficile* EGRIN model emerges as a resource to assign putative functions to uncharacterized genes based on functional associations among co-regulated genes (i.e., guilt-by-association)(Wolfe et al., 2005). We predicted putative functions for 48 uncharacterized genes by mining underlying functional enrichment of modules under different experimental conditions (**Table S4**; *see methods*). These 48 previously uncharacterized genes were associated with 13 functional categories, including “Sporulation” and “Other sugar-family transporters” (**Fig. 3A**).

**Figure 3.**
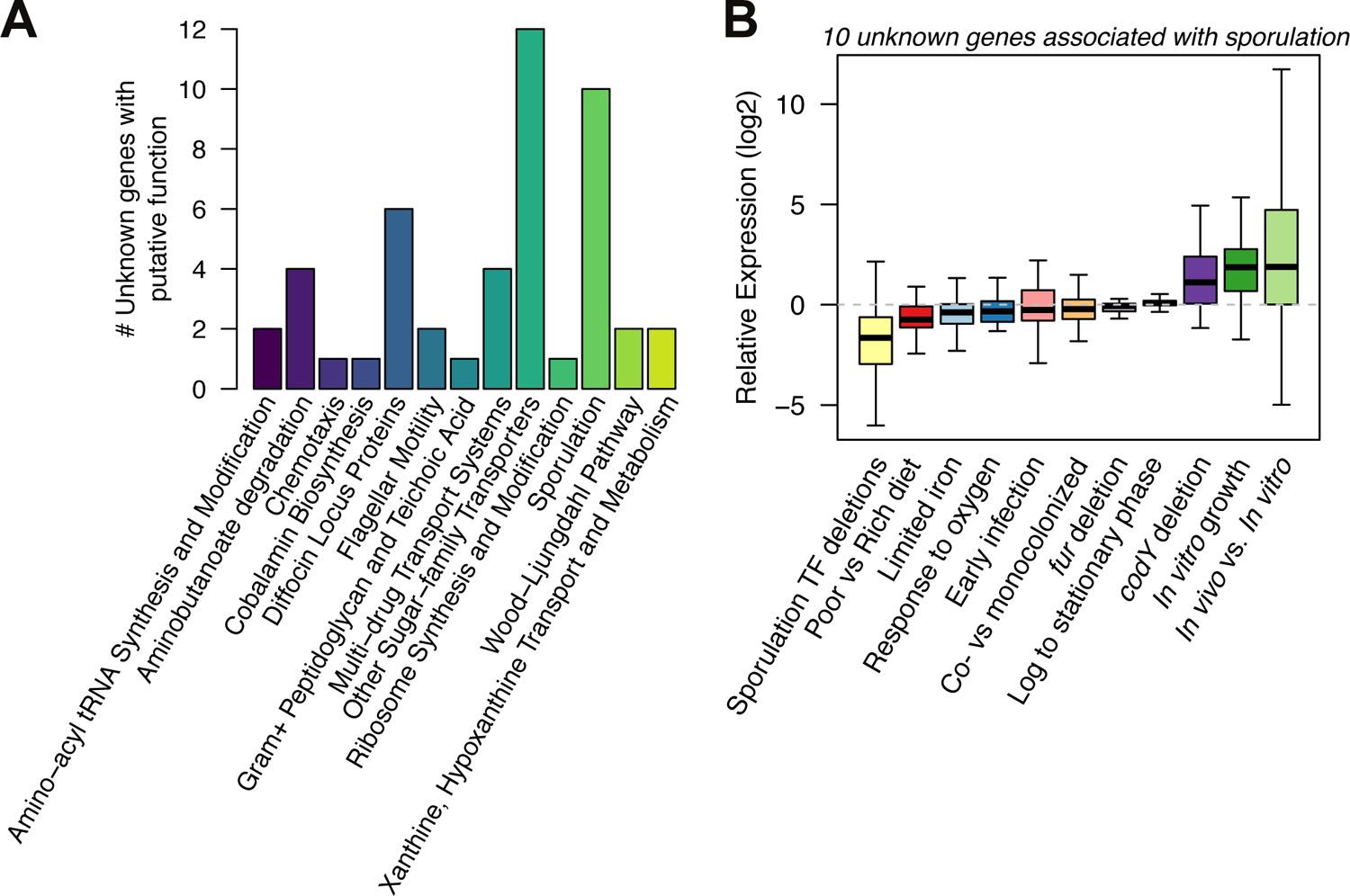
The EGRIN model offers insights on potential functions of uncharacterized genes of *C. difficile*. Hypotheses regarding the functions of 48 uncharacterized genes were generated based on their membership in high quality co-regulated gene modules significantly enriched with specific functional terms. (A) Barplot with the number of unknown genes associated with each functional term (from the *C. difficile* genome annotation in (Girinathan et al., 2021)). (B) The involvement of 10 uncharacterized genes in sporulation was supported by the observed strongest and significant downregulation in single deletion strains of transcriptional regulators of sporulation (*spo0A*, *sigEFGK*, *spoIIID*).

Ten genes were putatively assigned sporulation-related functions based on their co-regulation in the sporulation associated modules #206 and #251 (**Fig. 3B**). Module #206 includes seven stage III sporulation genes (*spoIIIAA*, *spoIIIAB*, *spoIIIAC*, *spoIIIAD*, *spoIIIAF*, *spoIIIAG* and *spoIIIAH*), and two stage IV sporulation genes (*spoIV*, *spoIVA*) (**Fig. S3A**). Module #251 includes the sporulation-associated alternative sigma factors SigG and SigE (located in the same operon) (**Fig. S3B**). Decreased expression of the 10 putative sporulation genes upon deletion of sporulation-associated sigma factors suggested putative roles within the mother cell or the forespore. Seven genes are likely associated with mother cell-specific roles based on their decreased expression in *sigE* (six genes) and *sigK* (one gene) deletion strains (**Table S2**). Two additional genes were downregulated in a *sigG* deletion strain, suggesting putative functions in the forespore. Notably, Tn-seq studies for gene essentiality in *C. difficile* identified seven of these 10 genes as required for sporulation (Dembek et al., 2015) (**Fig. S3A-B**).

Module #48 contains two adjacent operons (*4hbd*-*cat2*-CD630_23400-*abfD* and *sucD*-*cat1*) associated with aminobutanoate degradation (**Fig. S3C**). Both operons are regulated by CodY and PrdR. Hence, we predicted that the four uncharacterized genes in this module may be also involved in amino acid metabolism (**Table S4**). In support of this hypothesis, CD630_08760 and CD630_08780 are both differentially expressed upon *codY* deletion. Recent studies also suggest that CD630_ 08760 may function as a tyrosine transporter per its homology to the CodY-regulated neighbor gene, CD630_08730 (Bradshaw et al., 2019). A finding supported by the observed decreases in tyrosine uptake and Stickland fermentation in clinical isolates lacking CD630_08760 and CD630_08780 (Steglich et al., 2018). Altogether, we assigned 48 genes to putative functions in sporulation (10 genes), sugar transport (12 genes), among others. These examples illustrate how integration of multiple sources of evidence for context-specific co-regulation of genes in EGRIN can also predict putative functions for uncharacterized genes. EGRIN can also aid in designing experiments (e.g., perturbations of TFs and target genes) to validate predicted function assignments in relevant environmental context.

### EGRIN uncovers differentially active regulatory networks during *in vivo* infection

We investigated the differential expression of EGRIN modules across multiple *in vivo* studies (Janoir et al., 2013; Janvilisri et al., 2010) to discover the regulatory mechanisms that drive *C. difficile’s* colonization and adaption to *in vivo* environments. Analyses identified 680 genes across 43 modules that were significantly upregulated *in vivo*, while 1,325 genes across 82 modules were significantly downregulated. Notably, three modules were also differentially regulated during early stages of infection (**Supplementary File S1**). For example, this analysis discovered the *in vivo* activation and repression of module #158 and module #48, respectively (**Fig. 4A-B**). Notably, module #48 was upregulated during early infection and it is enriched with members of the CodY and PrdR regulons, as described above. Module #158 is enriched for putative PrdR and EutV co-regulated ethanolamine utilization genes, including *eut* operons for a two-component histidine kinase sensing system and carboxysome structural proteins that house the ethanolamine fermentative enzymes (Nawrocki et al., 2018) (**Fig. S3D**). Ethanolamine is prevalent within gut secretions and is also released from damaged host tissues, providing a readily available carbon and nitrogen source for *C. difficile*. The predicted co-regulation of this gene module by PrdR (Pearson correlation = −0.29 and P-value = 9.7e-04) suggests additional *in vivo* functions of this regulator to optimize *C. difficile’s* metabolism in gut environments. The negative relation between *prdR* expression and module #158 expression indicates that enhanced utilization of ethanolamine as it becomes a newly available carbon and nitrogen source *in vivo* may contribute to the decreased survival observed in hamsters infected with a *prdR* deletion strain (Bouillaut et al., 2019).

**Figure 4.**
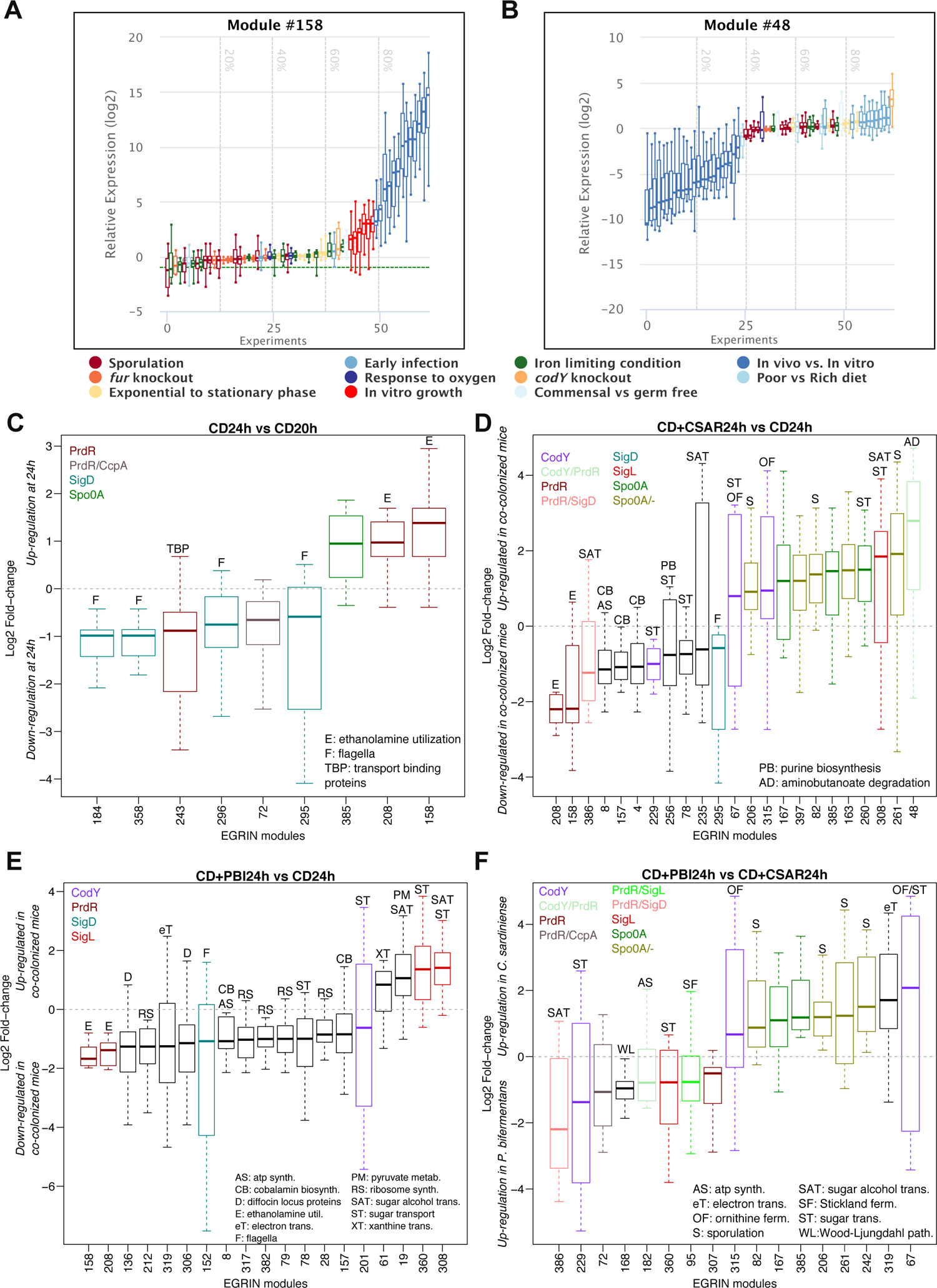
The EGRIN model identifies TFs driving the *in vivo* response of *C. difficile* when interacting with gut commensals *P. bifermentans* (PBI) and *C. sardiniense* (CSAR). (A) Expression profile of module #158 across the transcriptional compendium used to build the EGRIN model. Each box represents the log2 fold-change of all members of the module in a single transcriptome. Transcriptomes are color coded according to their membership in the 11 condition blocks included in the compendium. Transcriptomes were ranked based on their median log2 fold-change. Only one condition block (‘In vivo vs. in vitro’) was found in the 20% of highest (indicated with the 80% dashed line) fold-change. (B) Expression profile of module #48. The ‘Early infection’ condition block is statistically over-represented in the 20% of highest transcriptomes median log2 fold-change. (C) EGRIN modules enriched with genes differentially expressed (absolute log2 fold-change > 1 and adjusted p-value < 0.05) in *C. difficile* mono-colonized mice at 24 vs 20 hours of infection. X-axis shows module IDs. Modules were annotated according to their functional enrichment and overlap with manually curated TF regulons (Table S2). (D) Enriched EGRIN modules in *C. sardiniense*+*C. difficile* co-colonized mice vs *C. difficile* mono-colonized mice at 24 hours of infection. Due to space constraint, only abbreviations of functional terms not shown in other panels are displayed. (E) Enriched EGRIN modules in *P. bifermentans*+*C. difficile* co-colonized mice vs *C. difficile* mono-colonized mice at 24 hours of infection. (F) Enriched EGRIN modules in *P. bifermentans*+*C. difficile* co-colonized mice vs *C. sardiniense*+*C. difficile* co-colonized mice at 24 hours of infection. For all comparisons, only modules with absolute median fold-changes > 0.5, and enriched with TF regulons or functional categories are displayed.

With capacity to identify intestinal contributions to *C. difficile* responses, we leveraged the EGRIN model to analyze commensal modulation of the pathogen’s virulence, using transcriptomic datasets from gnotobiotic mice that were mono-colonized with the mouse-infective strain *C. difficile* ATCC43255 or co-colonized with *C. difficile* and the protective gut commensal species *Paraclostridium bifermentans* (PBI), or infection-worsening species *Clostridium sardiniense* (CSAR) (Girinathan et al., 2021). These datasets were not used in model construction. By mapping sets of differentially expressed genes into the EGRIN model we uncovered modules across 18 cellular processes and their associated TFs that were differentially regulated in the presence of PBI or CSAR (**Fig. 4C-F**).

One Spo0A-enriched module (module #385) was upregulated by 24h of infection in mono-colonized mice (**Fig. 4C**). The same module was upregulated by 24h of infection in CSAR co-colonized mice, in addition to three other modules enriched with sporulation genes (modules #82, #206, #261 in **Fig. 4D**). These three modules were also enriched with the Spo0A regulon. On the other hand, no sporulation-enriched modules were detected by 24h of infection in PBI co-colonized mice (**Fig. 4E**). Comparison of CSAR co-colonized mice and PBI co-colonized mice discovered four sporulation-enriched modules (modules #82, #206, #242 and #261) with increased expression in the virulent context (i.e., presence of CSAR) (**Fig. 4F**). These findings were confirmed with the high levels of spore release in expanded populations of vegetative *C. difficile* when co-colonized with CSAR (Girinathan et al., 2021). Overall, this analysis suggested that the sporulation pathway is an indicator of *C. difficile* disease, reinforcing the Spo0A-mediated link between sporulation and toxin production recapitulated by the model (**Fig. 2F**).

Module #319 contains multiple genes associated with electron transport via Rnf ferredoxin systems, and steps in glycolytic, butanoate and succinate metabolic pathways (**Fig. S3E**). This module was consistently downregulated in mice co-colonized with the protective commensal PBI (**Fig. 4E**), and was upregulated in CSAR co-colonized mice when compared to PBI co-colonized mice (**Fig. 4F**). These findings highlight activation of multiple co-regulated energy generating pathways in hypervirulent states of *C. difficile* which may contribute to virulence and to the pathogen’s responses to changing redox states brought on by severe host inflammatory responses. Remarkably, the EGRIN model identified the NAD+/NADH sensing regulator Rex as a potential repressor of module #319 (Inferelator regression coefficient = −0.075). The observed downregulation of module #319 in PBI co-colonized mice indicates increased Rex activity when *C. difficile* was introduced with this commensal and associated nutrient depleted state to support *C. difficile* growth (Girinathan et al., 2021). In support of a positive influence of PBI on Rex activity, we observed downregulation (suggesting decreased activity) and upregulation of *rex* mRNA level in the compiled transcriptional compendium during *in vitro* growth and early infection, respectively (**Fig. S3F**).

Four modules enriched with the SigD-regulated genes encoding subunits of flagella (modules #184, #295, #296 and #358) were downregulated in mono-colonized mice at 24h (**Fig. 4C**), indicating repression of motility to divert cellular resources toward pathogenesis. This finding is supported by increased virulence of *C. difficile* strains lacking a functional flagella (Dingle et al., 2011). In summary, the analysis of differential regulation patterns of EGRIN modules has generated new mechanistic insights and hypotheses for the interplay of specific genes and TFs for sporulation, energy production, and flagella synthesis that might underlie the enhanced and subdued virulence of *C. difficile* in different contexts.

### Metabolic network analyses elucidate *in vivo* metabolic adaptations of *C. difficile*

To investigate how specific genes within *C. difficile* contribute to *in vivo* phenotypes needed to develop symptomatic infection we extended a previously developed icdf834 metabolic model for CD630 (Kashaf et al., 2017; Larocque et al., 2014). We added four genes (CD630_08700, CD630_08680, CD630_17090 and CD630_10810) that encode reactions for molybdenum utilization and cofactor synthesis, and added exchange reactions to account for *C. difficile’s* capacity to utilize mannitol, fructose, sorbitol, raffinose, succinate and butanoate (Janoir et al., 2013; Theriot et al., 2014). In total, we added 46 reactions. We refer to this updated model as icdf838 (**Fig. 5A**). Lastly, the model derived from CD630 was compared with the gene feature content from *C. difficile* ATCC43255, used as a model organism to investigate symptomatic infections in mice. The two strains shared 92% of metabolic genes and predicted pathways (768 out of 838 genes in the icdf838 model have homology with the ATCC43255 strain; **Supplementary File S2**).

**Figure 5.**
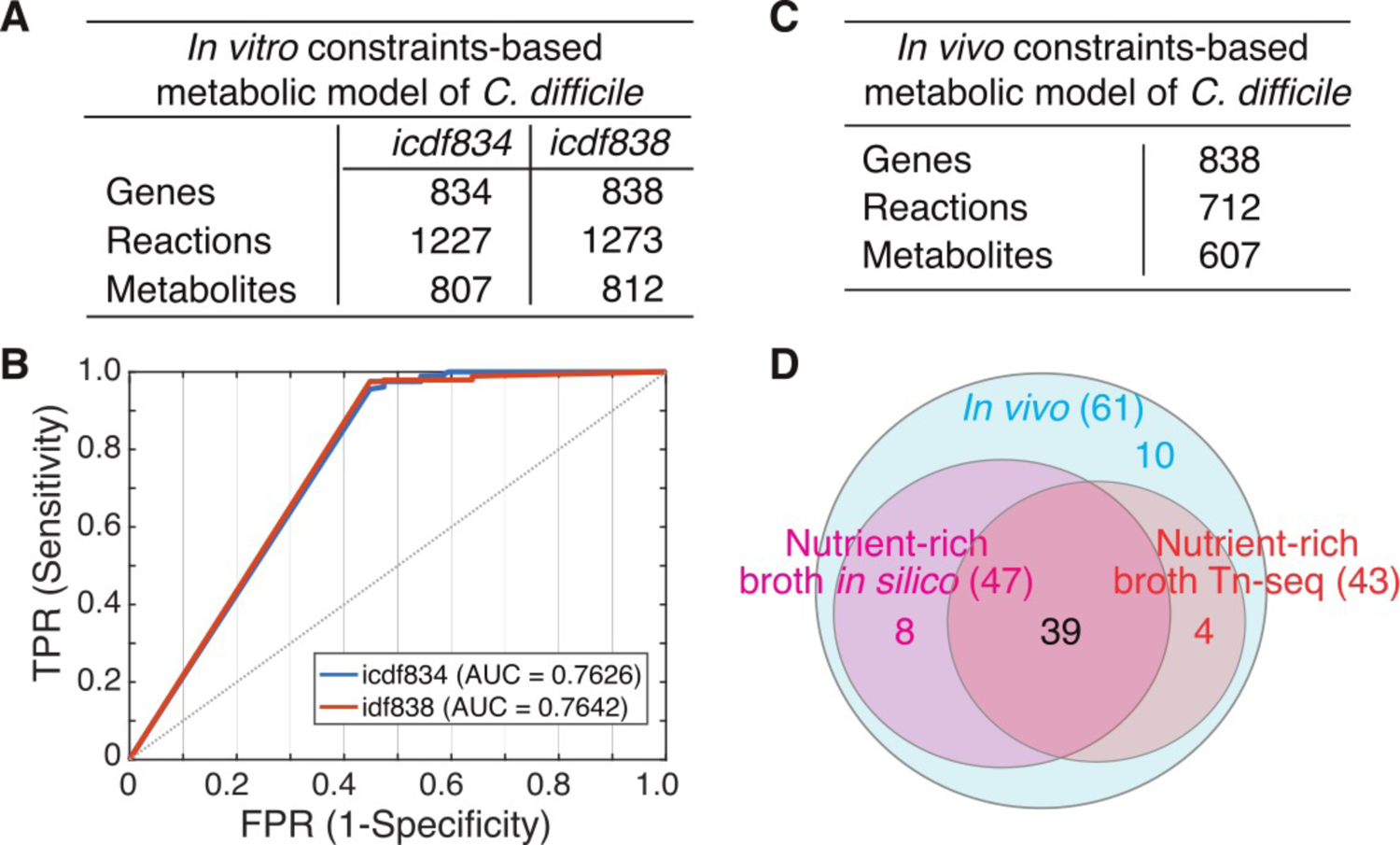
Metabolic model predictions. (A) Details of the *in vitro* metabolic models (icdf834 and icdf838) of *C. difficile* 630 (Kashaf et al., 2017). The number of genes, reactions and metabolites in the icdf838 model (generated in this study) after adding the required *in vivo* exchanges, transports and new reactions are shown. (B) ROC curves showing the accuracy of icdf834- and icdf838-predicted gene essentiality in nutrient rich medium evaluated against a Tn-seq functional screen (Dembek et al., 2015). (C) Details of the *in vivo* model derived using the GIMME algorithm (Becker and Palsson, 2008) on the icdf838 model where only the active reactions are included from *in vivo* transcriptome. (D) Venn diagram showing the number of model-predicted essential genes for growth of *C. difficile* 630 *in vitro* vs *in vivo* (i.e., mono-colonized condition).

We validated the completeness and accuracy of this model by confirming its ability to predict biomass production in three different *in vitro* media compositions: 1) minimal medium, 2) basal defined medium and 3) complex, nutrient-rich medium (see Larocque et al., 2014 for media compositions). The model accurately predicted *C. difficile’s* requirements for six amino acids: cysteine, leucine, isoleucine, proline, tryptophan and valine (Karasawa et al., 1995). Using a Tn-seq generated genome-wide fitness screen (Dembek et al., 2015), we tested the accuracy of model-predicted importance of each gene in supporting the growth of *C. difficile* in nutrient rich conditions using a receiver-operator characteristic (ROC) curve. The area under the ROC curves for the icdf834 and icdf838 models were 0.763 (p-value = 0.015) and 0.764 (p-value =0.029), respectively (**Fig. 5B**), indicating that both models significantly outperformed a random model.

We next extended and applied the model to predict *C. difficile* behaviors and gene essentiality *in vivo. C. difficile* transcriptomes from specifically-colonized gnotobiotic mice (Girinathan et al., 2021) were used as input into the GIMME algorithm (Becker and Palsson, 2008). Per the *in vivo* gene expression, we determined that 712 of the 1,273 total reactions in the icdf838 model were active and likely to be important for colonization and growth over the course of symptomatic infection (**Fig. 5C**). Leveraging information from the *in vitro* studies, the model made two notable predictions *in vivo* regarding the pathogen’s metabolism. First, the icdf838 model predicted 15 amino acids to be required for *C. difficile* growth *in vivo*, in contrast to the 6 required *in vitro* (**Supplementary File S2**). These amino acids included the dominant Stickland-fermented amino acids that were also required *in vitro,* including proline and branched chain amino acids, and additional amino acids including arginine, glutamate, lysine and methionine, which also have multiple cellular functions in cell wall synthesis, nitrogen cycling, and responses to oxidative stress. Secondly, the model predicted *C. difficile’s* switch from preferential use of glucose as a carbon source *in vitro* in complex media, to simultaneous utilization *in vivo* of diverse carbohydrate sources including fructose, galactose, maltose, and sugar alcohols such as mannitol and sorbitol, to promote colonization and growth (**Supplementary File S2**). Seven of these carbohydrate sources were described in other *in vivo* mouse infection studies illustrating support for these findings across *C. difficile* strains, and in germfree and conventional mouse models (**Supplementary File S2**) (Janoir et al., 2013; Jenior et al., 2017; Theriot et al., 2014).

### Integration of EGRIN and metabolic model reveals how the interplay of regulation and metabolism governs *C. difficile* adaptations to *in vivo* host environments

To evaluate how transcriptional regulation modulates *C. difficile*’s metabolic and physiological responses, we integrated the transcriptional and metabolic networks into a **P**henotype of **R**egulatory influences **I**ntegrated with **M**etabolism and **E**nvironment (**PRIME**) model (Immanuel et al., 2021). The PRIME model evaluates context-specific TF essentiality, an approach used to uncover vulnerabilities of *M. tuberculosis* (Immanuel et al., 2021). To generate a PRIME model for *C. difficile*, we first inferred a transcriptional network of 1,405 TF-gene interactions involving 215 TFs and 733 metabolic genes. This step was necessary to quantify combinatorial and conditional influence of TFs in metabolic genes and their associated metabolic fluxes, and to compensate for the poor coverage and lack of information about TF-gene interactions in *C. difficile*. Condition-specific PRIME models were generated by integrating *C. difficile* transcriptomic data from mice mono- and co-colonized (*C. difficile* and PBI), and using metabolic constraints for *in vitro* growth.

PRIME predicted essential metabolic genes and networks that promote *C. difficile’s* growth *in vivo* (i.e., in mono-colonized mice). In simulating the consequences of single gene deletions, PRIME predicted that *in vivo* growth of the pathogen in mono-colonized mouse would decrease by ≥65% with individual knockouts of 10 genes involved in one carbon-cycling reactions, and nucleotide and central carbohydrate metabolism (**Fig. 5D, Fig. 6A** and **Table S5**). These metabolic pathways represent new potential targets that drive *C. difficile’s* colonization and subsequent growth, factors required to develop symptomatic infections. Model predictions also illustrated *C. difficile’s* predicted shift from carbohydrate utilization towards amino acid utilizing pathways *in vivo*, as shown by the enhanced set of 15 amino acids, including the preferred Stickland donor and acceptor amino acids (leucine and proline) known to support metabolism and growth (Janoir et al., 2013; Jenior et al., 2017; Theriot et al., 2014). Notably, many of these amino acids show high abundance within the gut lumen in gnotobiotic and in antibiotic-treated conventional mice that enhance *C. difficile’s* capacity to colonize and expand (Girinathan et al., 2021).

**Figure 6.**
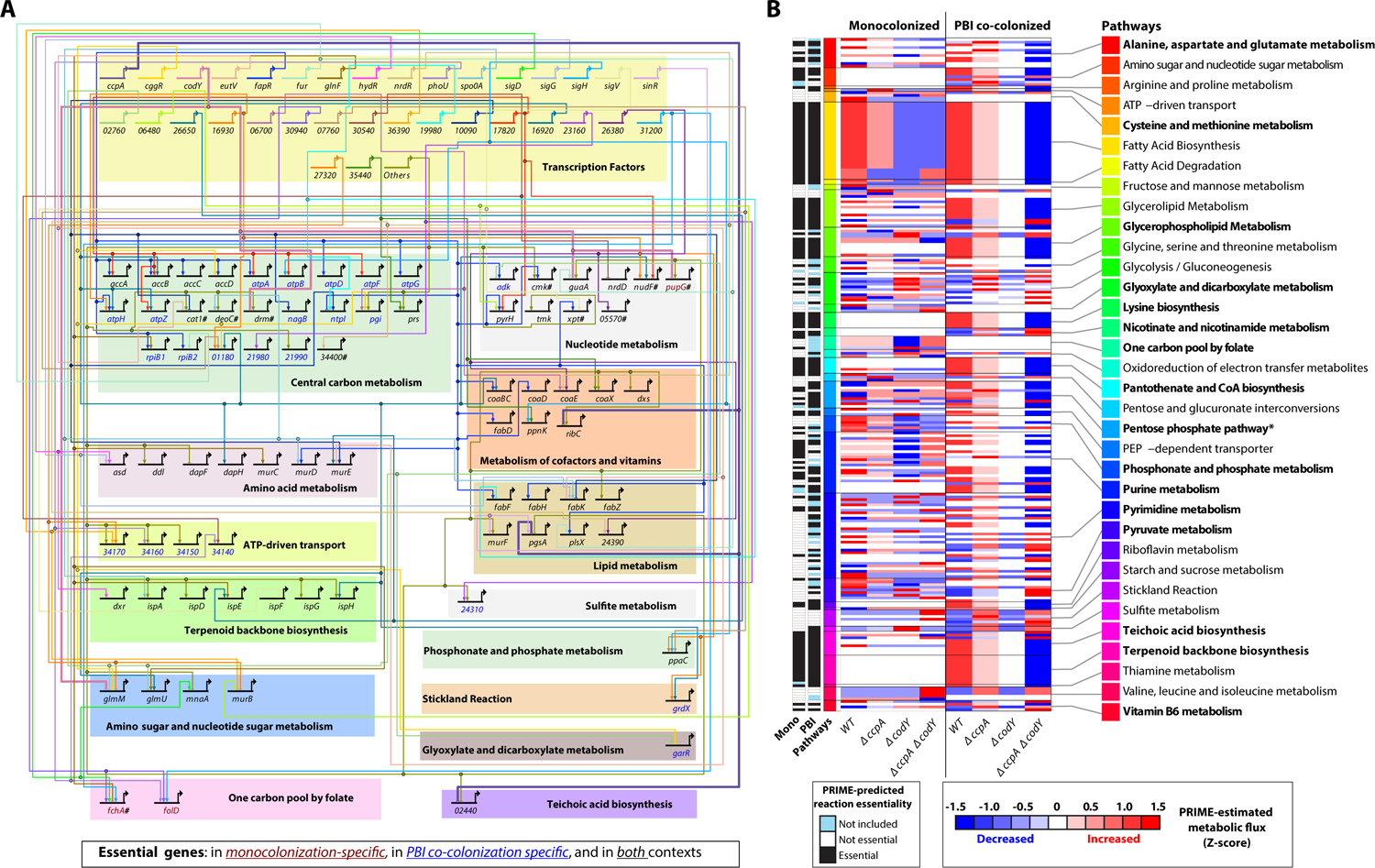
PRIME-estimated metabolic fluxes and gene essentiality in mono- and PBI co-colonized conditions. (A) BioTapestry visualization of *in vivo* gene regulatory network for *C. difficile* 630: 79 essential genes with predicted transcriptional regulators are shown. Transcriptional interactions were predicted using the Inferelator. The genes and regulators shown as five digit numbers represent the nomenclature preceded by ‘CD630_’. All regulators with less than three essential targets were combined in the ‘Others’ meta-regulator. The ‘#’ symbol indicates the 10 genes that are essential in the mono-colonized condition but not essential for *in vitro* growth. (B) PRIME-estimated metabolic fluxes for each reaction in the mono- and co-colonized conditions. Z-score transformation was independently applied to each reaction and condition. The ‘*’ symbol indicates that there is some contention about the existence of this pathway in *C. difficile* and it may need to be updated in future model revisions.

One of the key advantages of PRIME is its extended capability to evaluate TF essentiality. We evaluated the performance of PRIME by generating ROC curves as described above. The area under the ROC curve for PRIME gene essentiality predictions was comparable with corresponding values for the icdf834 and icdf838 models (**Fig. S4**). Given the influence of CcpA and CodY on toxin production and regulation of multiple metabolic genes (Antunes et al., 2012; Dineen et al., 2010), we leveraged PRIME to predict the phenotypic consequences of single and double knockouts in *ccpA* and *codY* in both mono- and PBI co-colonized mouse infections (**Table 1**). Using condition-specific models, PRIME predicted that three genes regulated by CcpA (*pgsA*, *ribC* and CD630_02440) were essential in both mono- and co-colonized conditions (**Fig. 6A**). Similarly, PRIME predicted that three genes regulated by CodY (*drm*, *glmM*, *guaA*) were essential in mono- and co-colonized mouse and one additional gene (*pupG*) was only essential in PBI co-colonized mouse (**Fig. 6A**). Strikingly, PRIME predicted that a double knockout of CcpA and CodY would have a synergistic consequence on inhibiting *C. difficile* biomass (as compared to the effects of individual knockouts in each TF), *especially* in PBI co-colonized mouse. Five out of the six PRIME essentiality predictions were validated by performing infections of germ free and PBI colonized mouse with wild type and mutant strains of *C. difficile* (**Table 1**). In addition, PRIME offered mechanistic insights into the conditional phenotypes of *C. difficile* by revealing how the single and double knockouts of *ccpA* and *codY* remodel the metabolic state of the pathogen in mono- and co-colonized contexts (**Fig. 6B**). For example, reactions associated with the one carbon pool by folate pathway were inactive in the co-colonized condition but were differentially affected by the gene knockouts in the mono-colonized mouse. PRIME also predicted that flux for reactions associated with teichoic acid biosynthesis and terpenoid backbone biosynthesis was invariable in the mono-colonized mice but sensitive to gene deletions in co-colonized mice. In summary, by providing mechanistic insights into how transcriptional regulation propagates throughout the metabolic network to manifest in fitness, PRIME offers a new capability to investigate how *C. difficile* adapts to complex biotic and abiotic changes within the *in vivo* gut environment.

### The C. difficile Web Portal, a resource for the *C. difficile* community

We have released a new *C. difficile* Web Portal (http://networks.systemsbiology.net/cdiff-portal/) to provide a discovery and collaboration gateway for the *C. difficile* scientific community. The portal aims to accelerate the advancement of the science and understanding of *C. difficile* biology, and the relationship between gene regulation, metabolism and virulence. Within the portal users can access publicly available datasets (e.g., transcriptional compendia), models, software and supporting resources. The Portal includes information on more than 4,000 *C. difficile* genes, 1,273 metabolic reactions, and 406 co-regulated gene modules. Notably, the EGRIN model can be interactively explored, and queried with gene set enrichment analysis to elucidate regulatory networks that may be relevant for a user-specified group of genes. Genes can be explored in the context of genome annotations, expression profiles, regulatory and metabolic membership, and other functional genomic information across different databases including COG, Uniprot, and PATRIC (Consortium, 2017; Galperin et al., 2015; Wattam et al., 2014). The portal provides access to detailed information on (1) Genes, (2) predicted Gene Modules, and (3) Metabolic Reactions (**Fig. S5A**).

Each gene module page includes summary statistics for the module, context-specific differential expression patterns of module genes, GREs, TF regulatory influences, enrichment of biological functions and pathways, and information on each regulon member gene. The module pages are structured to facilitate the assessment of the quality and statistical significance of the modules and highlight functional connections, while allowing users to implement their own filters (e.g., regression coefficient and adjusted p-value thresholds) (**Fig. S5B**). The portal includes a table of metabolic reactions with details of each reaction, associated genes, metabolites, and sub-systems. Metabolites and sub-systems are defined as taxonomic vocabularies that collect and group associated reactions to identify related metabolic processes. In addition, the portal provides access to algorithms, software, and data, and will include information about animal models, strains, and other *C. difficile* relevant community resources. As additional datasets are communicated, model predictions and tools will be successively enhanced to support systems-level analyses and assist in hypothesis generation in *C. difficile* biology and to enable tangible clinical interventions.

## DISCUSSION

The obligate anaerobe *C. difficile* is unique among gut anaerobes in possessing a diverse carbon source metabolism to enable colonization and growth in gut environments. These systems further exist within a complex network of gene regulatory modules that modulate growth, energy balance, and stress responses *in vivo*. Capacity to understand these systems-level integration points has remained challenging in the absence of robust systems biology models to infer *C. difficile’s in vivo* behaviors. We acknowledge the detailed studies from multiple groups over prior decades that provided a critical mass of information on *C. difficile’s* nutrient and gene-level responses to support development of an EGRIN model, the first for a gut anaerobe and toxigenic species. We emphasize that this information, the most for any obligate anaerobe, still represents a small fraction of that normally used to develop thorough EGRIN models. Recent improvements in the genetic manipulation of *C. difficile*, including the mouse infective strain ATCC43255 (Peltier et al., 2020), open new capacity to probe the GREs modulating critical aspects of its metabolism, growth and virulence, from a systems-level perspective.

The *C. difficile* EGRIN model enables a number of predictions relevant to *in vivo* disease. For example, the PrdR regulator of the pathogen’s Stickland proline reductase (*prdABDEF*) and other genes, has long been hypothesized to have a role in PaLoc gene expression through as-yet unknown mechanisms. EGRIN predictions included gene module #182, which identified combined PrdR and CodY effects on *tcdA* gene expression (suggested by the enrichment of module #182 with both PrdR and CodY regulons), providing a regulatory integration point and broader set of co-regulated genes to support further experimental analyses of co-regulation between these two TFs, including effects on PaLoc expression. Biclustering also identified interactions between Spo0A, another regulator hypothesized to modulate PaLoc expression, and *tcdB* expression in module #397. The identified modules, associated genes and regulators provide new information to support further experimental investigation of combinatorial effects of these and other regulators identified in PaLoc gene-associated modules. The EGRIN model also predicted PrdR as a critical regulator *in vivo* through its systems-level effects on critical metabolic and regulatory networks supporting the pathogen’s colonization, metabolism and growth. These interactions mediated by PrdR involve multiple direct and indirect effects upon other modules (e.g., module #48 and module #158) and aspects of the pathogen’s metabolism and gene regulation.

The current model did not identify all experimentally known regulators of PaLoc expression, including SigD regulation of TcdR, and effects of other more recently identified PaLoc regulators such as RstA and LexA, for which limited datasets exist from wild type or targeted deletion mutants under relevant nutritional and other environmental perturbations. Additional datasets with isogenic regulator mutants will likely improve predictions while further validating previously defined biologic effects. Nonetheless, as shown with our *in vivo* analyses, application of the EGRIN and PRIME models to new datasets offers key insights into causal mechanistic drivers of adaptive strategies of the pathogen. Given that less than 10% of transcriptomic information and less than 2% of ChIP-seq regulator datasets were available for CD630, as compared to EGRIN models developed for other species, the model provides a formative tool to design future transcriptomic and ChIP-seq studies to improve predictions for these regulons.

Leveraging additional Tn-seq and *in vivo* transcriptomic datasets, the expanded icdf838 model identified a broader set of amino acids, in addition to genes and anaerobe-specific pathways, needed to support colonization and growth expansion *in vivo*. Notably, predictions of *in vivo* gene essentially identified folate one-carbon cycling pathways including those connected with Wood-Ljundahl fixation of carbon dioxide to acetate (Gößner et al., 2008). Predictions of gene essentiality also identified multiple nucleotide synthesis and salvage pathway genes that were essential *in vivo* but not *in vitro,* including ones associated with xanthine transport and metabolism, an abundant nucleotide in gut secretions that originates from host sources (Girinathan et al., 2021). Lastly, the systems approach using the two models has identified known and novel genes and TFs across disparate pathways, including sporulation, flagella biosynthesis, sugar metabolism, and biosynthesis of aromatic and branched-chain amino acids, that contribute to growth in an intestinal environment. Once experimentally validated, these essential genes represent novel vulnerabilities that can be rationally targeted with small molecules, bacteriotherapeutics, or other patient interventions. We believe that by democratizing the disparate data, algorithms, and models through interactive exploration capabilities, the *C. difficile* Web Portal will accelerate collaborative systems analysis of host-pathogen-commensal interactions by engaging the wider scientific community.

We illustrate additional predictions from the *C. difficile* EGRIN model to enable gene-through systems-level analyses of the pathogen. Though among the best described obligate anaerobes, the *C. difficile* genome still contains a high number of genes of unknown function. Model predictions provided new information to assign putative functions to 48 genes, including ones associated with sporulation, carbohydrate transport, and other aspects of cellular metabolism. Integration of the regulatory and metabolic network models with PRIME enabled unprecedented insight into the essential role of TFs in mediating combinatorial control of metabolism in different colonization contexts, vis-a-vis presence and absence of the protective commensal PBI. EGRIN and PRIME models will be especially powerful to navigate the vast space of possible combinatorial perturbations with *in silico* simulations and prioritize experiments that are most likely to reveal mechanistic insights into phenotypes that emerge from the complex interplay of regulatory and metabolic processes. For instance, PRIME accurately predicted enhanced synergistic epistasis between the CodY and CcpA networks, a phenotype that was not attributable to a single downstream metabolic gene, reaction or process. Rather, the significantly diminished growth of a double knockout of CodY and CcpA (especially in mice co-colonized with PBI) appears to emerge from the interplay of more than 35 TFs regulating ∼80 genes catalyzing reactions across more than two dozen metabolic processes. Importantly, these PRIME-derived biological insights can iteratively drive hypothesis-driven characterization of mechanisms by which the pathogen infects and colonizes a complex host environment. Furthermore, by uncovering conditionally essential genes and pathways, PRIME can be used to understand how *C. difficile* escapes therapies, and develop strategies to block these escape routes with synergistically acting secondary antibiotics. The *C. difficile* Web Portal makes all of the tools and resources available to the broader research community, providing a platform for collaboration and to support systems-level investigations of the pathogen and its interactions with the host and commensal microbiota.

## STAR METHODS

### *C. difficile* genome annotation

A new ATCC43255 reference genome was generated and annotated to support *in vivo* transcriptome studies of *C. difficile* per discrepancies noted in the RefSeq genome, particularly among bacteriophage loci and other mobile elements (Girinathan et al., 2021). The updated reference genome was annotated using the NCBI Prokaryotic Genome Automatic Annotation Pipeline (Tatusova et al., 2016), PATRIC (Wattam et al., 2014), and PROKKA (Seemann, 2014) to extract gene features for support of transcriptome pathway enrichment analyses. Bacteriophage loci and genes were identified using PHASTER (Arndt et al., 2016).

### *C. difficile* transcriptional compendium

To generate a transcriptional compendium for *C. difficile*, required for constructing an EGRIN model, a total of 151 publicly available transcriptomes of *C. difficile* 630 (identified by searching for the ‘Clostridioides difficile 630’ term) were downloaded from the NCBI Gene Expression Omnibus (GEO) repository (Barrett et al., 2012) in March 2020. Downloaded transcriptomes were generated by 11 independent studies (**Table S1**). To integrate this data into a single dataset, we computed the log2 fold-change of each transcriptome with respect to a control condition, as performed in the generation of other transcriptional compendia (Moretto et al., 2016; Peterson et al., 2014). This step was not necessary for transcriptional data collected with dual channel arrays that included a normalizing control channel. The resulting transcriptional compendium contained a total of 4,091 gene features and 127 conditions. The 127 conditions in the transcriptional compendium were organized in 11 distinct conditional blocks (e.g., sporulation, *fur* deletion), as shown in **Table S1**.

### Construction of the EGRIN model

The EGRIN model for *C. difficile* was constructed in two stages. First, we used cMonkey2 (Reiss et al., 2015), a biclustering algorithm, on the compiled compendium of 127 *C. difficile* transcriptomes to simultaneously detect co-regulated gene modules and the conditions where co-regulation occurs. cMonkey2 integrates functional annotation from the STRING database (Szklarczyk et al., 2016), gene promoter sequences from the RSAT database (Nguyen et al., 2018), and operon predictions from MicrobesOnline (Dehal et al., 2010) when detecting gene modules. cMonkey2 was run using default parameters. Briefly, we used 2,000 iterations to optimize the co-regulated gene modules, each one with 3-70 genes. In each iteration, cMonkey2 refined the gene modules by evaluating and modifying (if necessary) condition and gene memberships. cMonkey2 biclustering approach allowed genes and conditions to be assigned to a maximum of two and 204 different modules, respectively. *De novo* GRE search was performed using MEME v. 4.12.0 (Bailey et al., 2015). Second, we used the Inferelator (Arrieta-Ortiz et al., 2015), a network inference algorithm, to identify potential transcriptional regulators for the 406 gene modules generated by cMonkey2. The Inferelator uses a Bayesian Best Subset Regression to estimate the magnitude and sign (activation or repression) of potential interactions between TFs and gene modules. We bootstrapped the expression data (100 times) to avoid regression overfitting (Arrieta-Ortiz et al., 2015). The Inferelator generates two scores for each TF-module interaction, the corresponding regression coefficient (i.e., beta) and a confidence score. The second score indicates the likelihood of the interaction. The final set of TF-module interactions was defined as the 805 interactions with absolute beta values ≥ 0.1.

### Experimentally supported literature derived TF regulons

We mined available literature to compile a list of experimentally supported targets for the 13 partially characterized transcriptional regulators (involved in sporulation, motility, carbon metabolism, among other processes) shown on **Table S2**. The manually compiled regulons represented a total of 1,349 regulatory interactions and involved 1,044 genes. Target genes included in the compiled TF regulons were supported by transcriptional data, protein-DNA binding data and *in silico* analysis of promoter regions (e.g., presence of known regulators DNA binding motif).

### Module enrichment evaluation

We used a hypergeometric test to identify modules of co-regulated genes in the EGRIN model that were statistically enriched with manually compiled TF regulons (**Table S2**) or functional pathways derived from curated annotation of *C. difficile* genome (Girinathan et al., 2021). Only gene modules with adjusted hypergeometric test p-value ≤ 0.05 and containing five or more genes from the relevant TF regulon or functional pathway were considered enriched.

### Assignment of putative functions to uncharacterized genes

To predict the potential role of uncharacterized genes, gene functions were predicted based on the functional enrichment of EGRIN modules (evaluated as explained above). Function assignments were restricted to uncharacterized genes located in functionally enriched modules in which 45% of more of the annotated members (i.e., genes with putative function) were assigned to the over-represented function term. This second filter was implemented to focus on EGRIN modules that were involved in a specific function. Thus, increasing the likelihood that the uncharacterized genes were also involved in the same function.

### Analysis of *in vivo* data

*In vivo* transcriptomic data from gnotobiotic mice mono-colonized with *C. difficile* ATCC43255 or co-colonized with *P. bifermentans* or *C. sardiniense* (Girinathan et al., 2021) were analyzed as previously described using the updated reference genome of ATCC43255 to extract gene features for subsequent analysis with DESeq2 (Love et al., 2014).

### Metabolic model refinement and gene essentiality prediction

A published genome-scale metabolic model of *C. difficile* 630 strain, icdf834 (Kashaf et al., 2017), was used in this study and expanded by adding reactions required for *in vivo* survival of the pathogen. The icdf834 model incorporates 1,227 metabolic reactions and 807 metabolites. The metabolic reactions were mapped through gene-protein-reaction associations to 834 genes, which represent 80% of 1,030 annotated metabolic genes in the CD630 genome (**Fig. 5A**). We also curated pathway annotations that were incorrectly designated using default KEGG annotations (Kanehisa et al., 2017). For example, most anaerobes do not utilize the tricarboxylic citric acid (TCA) cycle, although some reactions, in reverse, support aspects of pyruvate, succinate and oxaloacete metabolism. In the icdf834 model, we changed subsystem pathway annotation of two reactions - i) acetyl-CoA:oxaloacetate C-acetyltransferase and ii) succinyl-CoA synthase from TCA cycle to pyruvate metabolism and butanoate fermentation respectively (**Supplementary File S2**). Similarly, we updated reactions originally assigned to gluconeogenesis and the pentose phosphate pathway. We evaluated the homology of metabolic genes between *C. difficile* 630 and ATCC43255 strain of *C. difficile* in order to use the icdf834 model for representing the *in vivo* infection state of ATCC43255 strain. The details of 768 genes that are predicted in this homology analysis is provided in **Supplementary File S2**. We then extended the model by including four new genes and eight new exchange reactions that are required for the growth of the pathogen in the *in vivo* micro-environment, based on KEGG annotations. We also expanded the proline reductase (PR) and glycine reductase (GR) systems by adding alanine, branched chain amino acids (valine, leucine and isoleucine) and aromatic amino acids (phenyl alanine, tyrosine and tryptophan) as donors in the Stickland metabolism (Jackson et al., 2006), increasing the total number of reactions from 1,227 to 1,273 (**Supplementary File S2**). We named this expanded version of the model as “icdf838”. Then, the transcriptome of *C. difficile* profiled from *in vivo* infections of specifically-colonized gnotobiotic mice (Girinathan et al., 2021) was mapped onto the icdf838 model using the GIMME algorithm (Becker and Palsson, 2008).

This resulted in a model with 712 active reactions, with no changes in the number of genes. This model represents the *in vivo* state of *C. difficile*. We applied the constraint-based method for simulating the metabolic steady-state of *C. difficile* using flux-balance analysis (FBA) (Becker and Palsson, 2008; Orth et al., 2010). The initial validation steps involved checking the capacity of the icdf834 model to produce biomass in defined media conditions including 1) minimal medium, 2) basal defined medium and 3) complex, nutrient-rich medium (compositions used according to Larocque et al., 2014). Then, we tested the performance of both icdf834 and icdf838 models using gene essentiality predictions by FBA. A gene was considered “essential” if its deletion reduced the biomass by >65%. By this analysis, the model classified each gene as “essential” or “non-essential”. We compared the gene essentiality predictions from nutrient-rich media constraints with the available experimental Tn-seq data (Dembek et al., 2015) and deduced the confusion matrix to derive true positive rates (TPR) and false positive rates (FPR). This led to the elucidation of sensitivity and specificity of the model using ROC curve analysis. We then applied the same strategy and predicted the essential genes *in vivo* using FBA with the expanded and GIMME-derived context-specific network, icdf838. All model simulations related to FBA were performed on MATLAB_R2019a platform using the recent version of COBRA (The COnstraint-Based Reconstruction and Analysis) toolbox (Heirendt et al., 2019). *In silico* gene essentiality predictions were performed using the COBRA toolbox ‘single-gene-deletion’ function in MATLAB. The illustration of essential gene regulatory network *in vivo* was deduced using BioTapestry tool (Paquette et al., 2016).

### PRIME model development

Three PRIME models were constructed for *C. difficile* during *in vivo* (mono- and PBI co-colonized mice) and *in vitro* growth. Briefly, we first inferred a TF-gene transcriptional network using the Inferelator with incorporation of TF activity, estimated using the signed (positive or negative) TF-gene interaction network compiled for 13 TFs (**Table S2**), as we have previously done for other species (Arrieta-Ortiz et al., 2015, 2020). To infer a global network, 288 putative transcriptional regulators without known targets were also included as potential regulators. The inferred network included 5,801 TF-gene interactions with absolute regression coefficients ≥ 0.1. The transcriptional network was then integrated with the icdf838 metabolic model as explained in (Immanuel et al., 2021). Importantly, PRIME models were made context-specific by excluding metabolic reactions associated with lowly expressed genes, as explained above.

### C. difficile Web Portal

This portal utilizes the powerful build, search, collaboration, and visualization features of the Drupal content management system. With the two key features of modularity and extensibility, Drupal provides a slim, powerful core that can be readily extended through custom modules and easy-to-use collaborative tools to support information sharing. Based on these key features, we developed this content management system into a data management, analysis, and visualization framework to support *C. difficile* research.

Due to the complexity of the information provided by the genome and models, it is critical to provide a user-friendly and flexible search and filtering capabilities. By taking advantage of Drupal’s built-in search interface and implementing Apache Solr search, we created very powerful search capabilities that will query every information included in the portal database. Moreover, the search interface uses “facets” to allow users to explore a collection of information by applying multiple filters. This combination together with sorting enables users to start with broad searches and then quickly pinpoint specific information.

In order to provide a comprehensive functional genomics resource for the *C. difficile* community, genome annotations from several different sources were merged and imported into the C. difficile Portal. Curated genome annotations for *Clostridium difficile* strain 630 published by Monot et al. (Monot et al., 2011), were downloaded from MicroScope platform (Vallenet et al., 2017). Additional functional annotations were downloaded from PATRIC (Wattam et al., 2014) and Uniprot (Consortium, 2017) and merged with curated genome annotations. Overall, 4,018 genes were included in the Cdiff Portal. The *C. difficile* genome included 1,030 metabolic genes, 309 TFs, 270 small non-coding RNAs (sRNAs) (Soutourina et al., 2013), 87 tRNAs, 32 rRNAs and 17 miscellaneous RNAs (miscRNAs). The genome included 1,330 genes with unknown function. Furthermore, gene essentiality data from Dembek et al. (Dembek et al., 2015) was integrated with gene annotations.

## ACKNOWLEDGEMENTS

We thank members of the Baliga lab for critical discussions and feedback. Funding was provided by National Institute of Allergy and Infectious Diseases (R01AI128215 and R01AI141953), and National Science Foundation (BDI-1565166) to NSB, and the BWH Precision Medicine Institute, Hatch Family Foundation, R01AI153605 and P30-DK034854 to LB. BPG was supported by T32 HL007627. JNW was supported by the Intramural Research Program of the National Library of Medicine, National Institutes of Health. Funding was also provided by the Institut Pasteur and Université de Paris to BD and JP, and the Institut Universitaire de France to OS.

## COMPETING INTEREST

LB is an inventor on patents for *C. difficile* microbiota therapeutics. LB is an SAB member and holds stock in ParetoBio Inc. Remaining authors declare no competing interests.

## AUTHOR CONTRIBUTIONS

Conceptualization: MLAO, LB and NSB. Data curation: MLAO, SRCI, ST, BPG, JNW, ND, JP, OS, BD and LB. Formal analysis: MLAO and SRCI. Investigation: MLAO, SRCI, JP, BD, LB and NSB. Methodology: MLAO, SRCI, ST and WJW. Project Administration: MLAO and NSB. Resources: JP, BD, LB and NSB. Software: MLAO, ST, WJW. Supervision: NSB. Visualization: MLAO, SRCI, ST, WJW, NSB. Writing: MLAO, SRCI, ST, LB and NSB.

## DATA AND CODE AVAILABILITY

All datasets, models, software and supporting resources are available in the interactive and open access C. diff Web Portal (http://networks.systemsbiology.net/cdiff-portal/).

## SUPPLEMENTARY FIGURES

**Figure S1.**
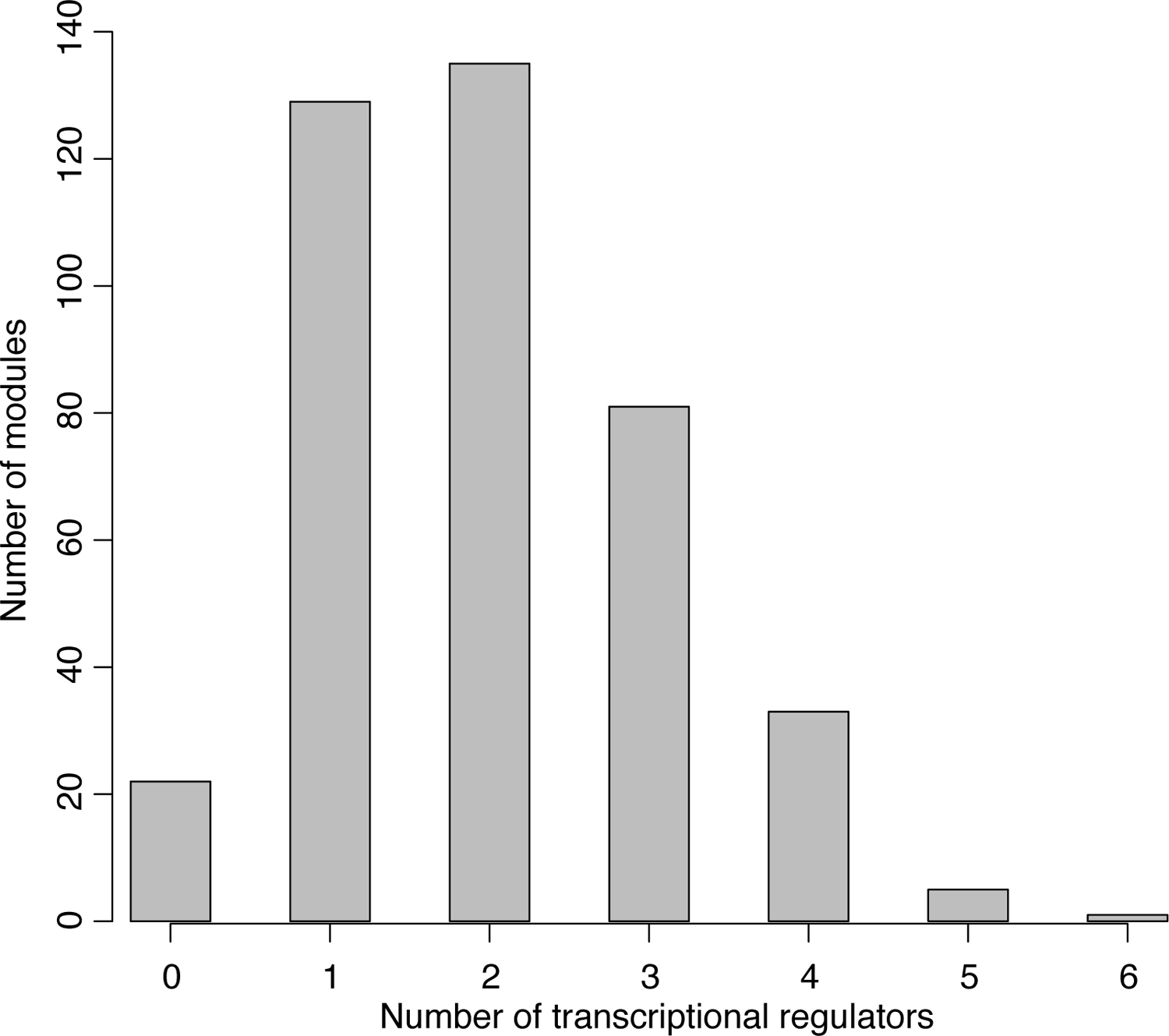
Number of Inferelator-predicted transcriptional regulators of gene modules in the EGRIN model.

**Figure S2.**
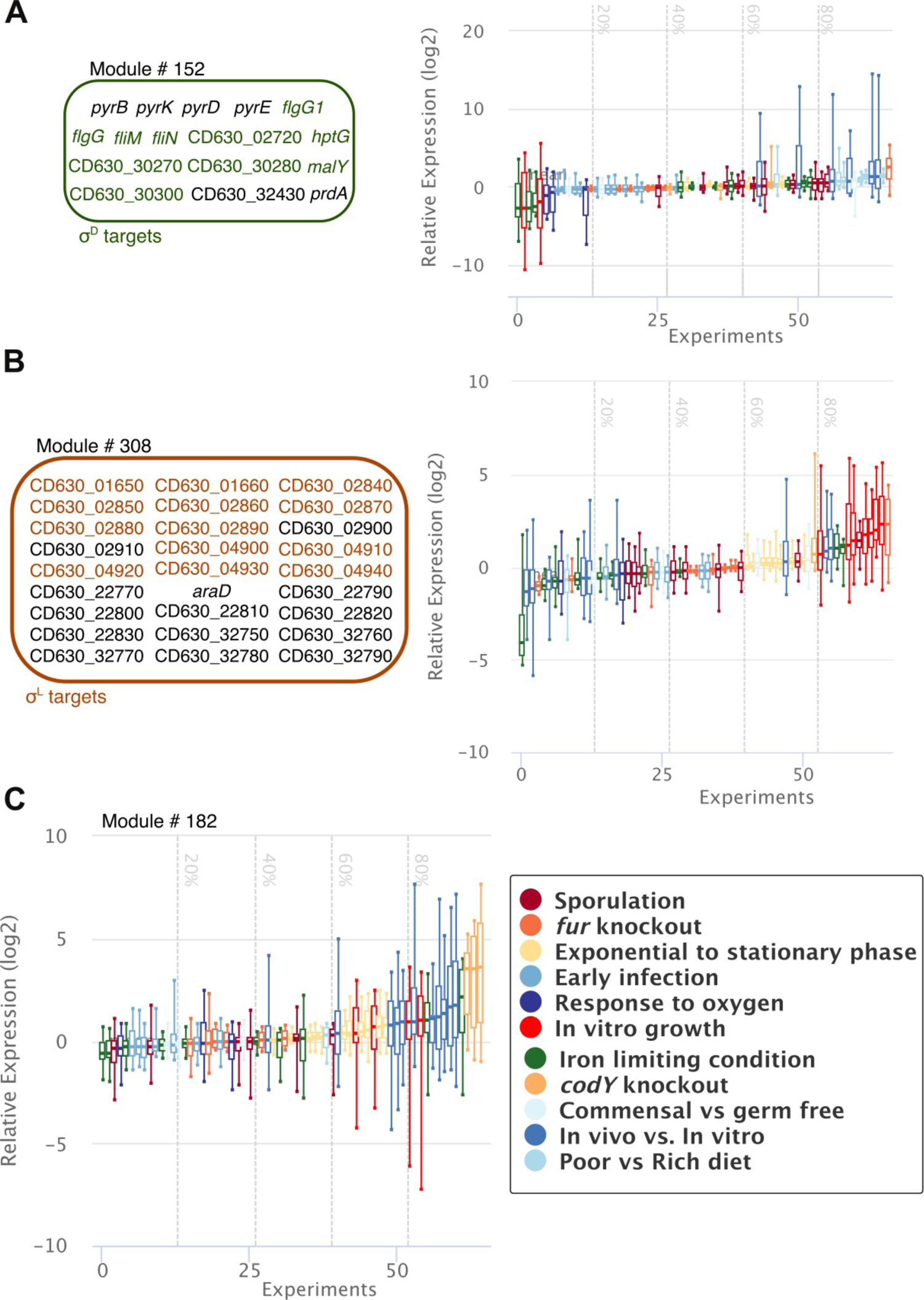
General properties of selected EGRIN modules. (A) Members of module # 152 are shown in the left panel. Transcriptional profile of module # 152 is shown in the right panel. Each box represents the log2 fold-change of all members of the module in a single transcriptome. Transcriptomes are color coded according to their membership in the 11 condition blocks included in the compendium (shown in the figure key). Transcriptomes were ranked based on their median log2 fold-change. (B) Members of module # 308 are shown in the left panel. Transcriptional profile of module # 308 is shown in the right panel. (C) Transcriptional profile of module # 182.

**Figure S3.**
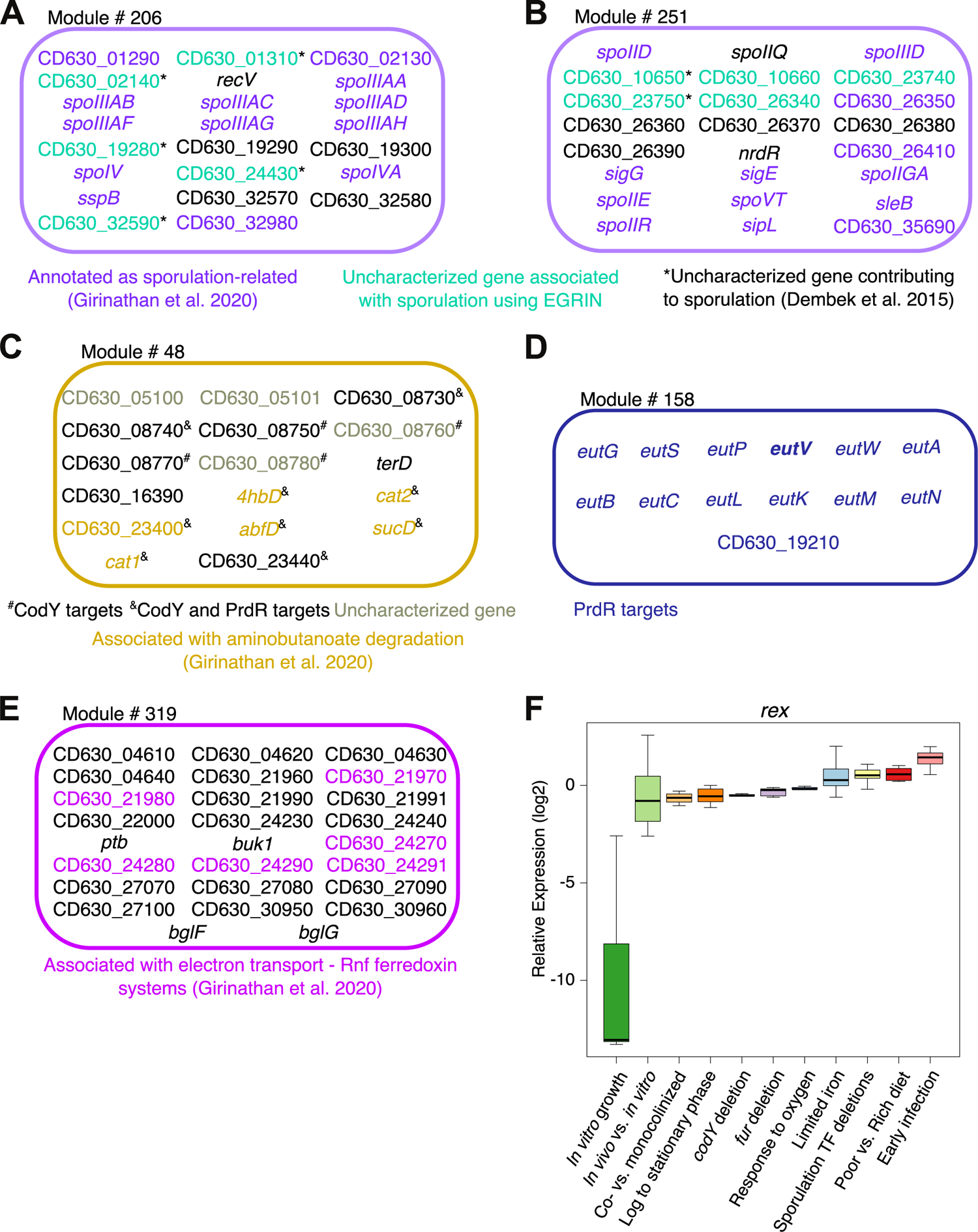
Modules associated with uncharacterized genes with putatively assigned function and *in vivo* adaptation. (A) Members of module # 206 are shown. (B) Members of module # 251 are shown. (C) Members of module # 48 are shown. (D) Members of module #158 are shown. (E) Members of module #319 are shown. (F) Transcriptional profile of *rex* in the transcriptional compendium compiled to build the EGRIN model.

**Figure S4.**
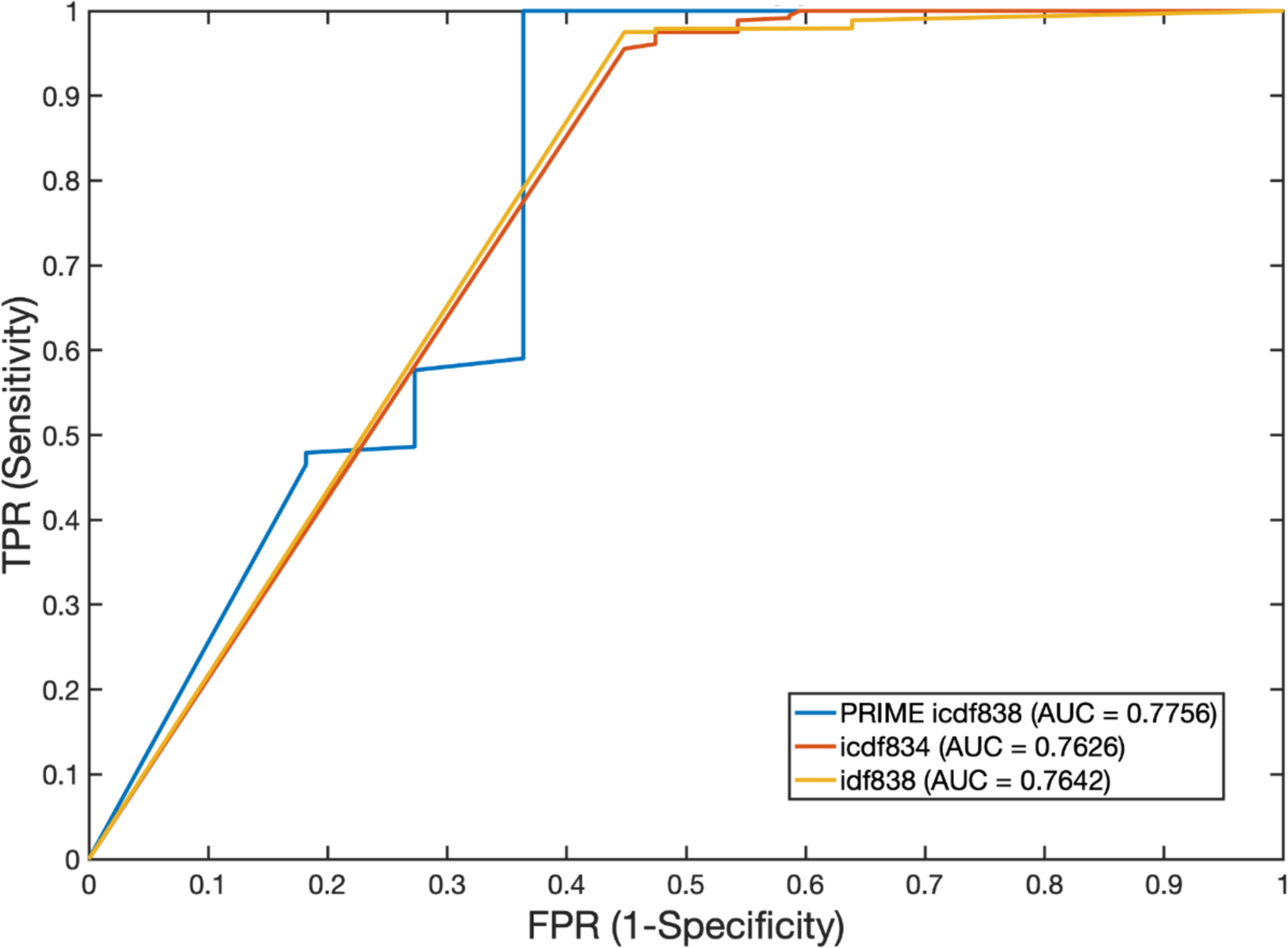
PRIME accurately predicts TF and gene essentiality. ROC curves showing the accuracy of icdf834-, icdf838 and PRIME icdf838-predicted gene essentiality in nutrient rich medium evaluated against a Tn-seq functional screen (Dembek et al., 2015). Area Under the Curves (AUC) values for all models are shown.

**Figure S5.**
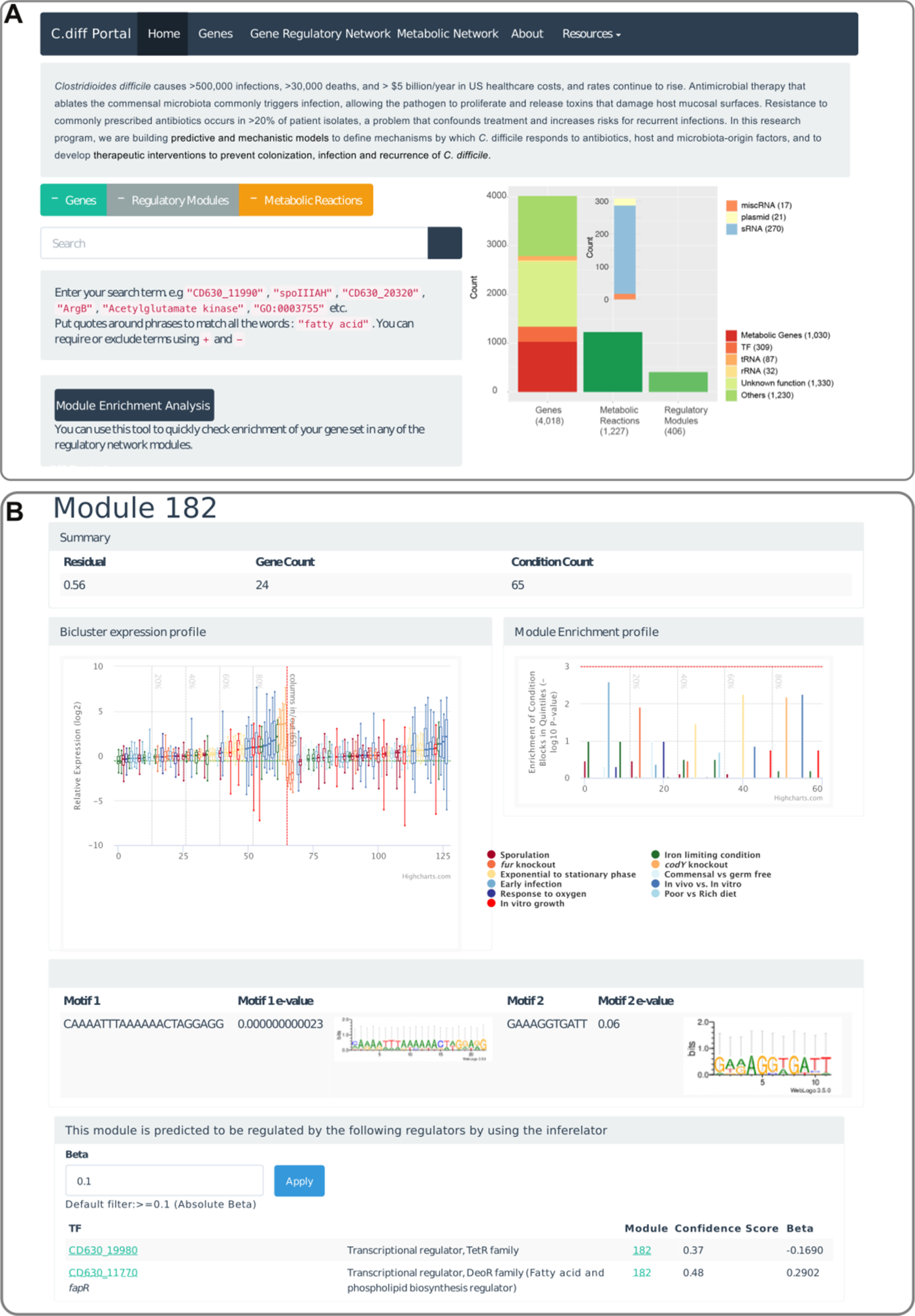
The Cdiff Web Portal. (A) Home page of the Cdiff Portal. Users can explore the gene regulatory network model and the metabolic model for *C. difficile*. In addition, all files are accessible in the resource tab. The search bar facilitates website exploration. (B) Module #182, associated with CodY and shown in Fig 2D is used as an example. Each module page includes general statistics of the module (residual score, gene count), displays the module expression profile in the compiled transcriptional compendium, the enrichment of each quintile of the module expression profile with the different condition blocks (e.g., early infection, etc.), and the detected GREs. A module page also offers information about the potential transcriptional regulators of the module. Putative regulators are defined based on over-representation of manually compiled TF regulons (assessed using hypergeometric test) and based on the Inferelator predictions.

## SUPPLEMENTARY TABLES

**Table S1.** Datasets used for generating *C. difficile* transcriptional compendium

| Condition | GEO Series Accession | # Transcriptomes<sup>a</sup> | # Controls<sup>b</sup> |
| --- | --- | --- | --- |
| Early infections<br>(0h, 30 mins, 60 mins, 120 mins) | GSE18407 | 12 | NA<sup>c</sup> |
| *In vivo* vs *in vitro*<br>(8h, 14h, 38h) | GSE43305 | 24 | NA<sup>c</sup> |
| *In vitro* growth<br>(8h, 14h, 38h) | GSE43305 | 8 | NA<sup>c</sup> |
| Iron limitation | GSE109453<br>GSE120189 | 15 | 15<sup>d</sup> |
| *fur* deletion | GSE69218<br>GSE120189 | 12 | 12<sup>d</sup> |
| Response to oxygen | GSE109175 | 3 | 3 |
| Response to commensals | GSE60751 | 8 | 8<sup>e</sup> |
| Poor diet vs Rich diet | GSE60751 | 8 | 8<sup>e</sup> |
| Transition from exponential to<br>stationary phase | GSE115054 | 16 | NA<sup>c</sup> |
| *codY* KO | GSE23192 | 3 | NA<sup>c</sup> |
| Sporulation TF deletions<br>(*spo0A* KO, *sigEFGK* KOs,<br>*spoIIID* KO) | GSE45977<br>GSE63777 | 18 | 6 |

<sup>a</sup>Refers to the number of arrays used as numerator when estimating log2 ratios
<sup>b</sup>Control arrays were averaged and used as denominator when estimating log2 ratios
<sup>c</sup>Not Applicable. Dual channel array and therefore the control was included in each array
<sup>d</sup>Six samples (half of the transcriptomes) in the GSE120189 dataset were used as controls to compute log2 ratios associated with the "Iron limitation" or "*fur* deletion" conditions
<sup>e</sup>Eight samples (half of the transcriptomes) in the GSE60751 dataset were used as controls to either compute log2 ratios for the "Response to commensals" condition or the "Rich diet vs Poor diet" condition

**Table S2.** Compiled TF regulons

| Regulator | Regulon Size | Supporting Data | Reference |
| --- | --- | --- | --- |
| CcpA | 194<sup>a</sup> | Protein-DNA binding, transcriptomics, *in silico*<sup>b</sup> | (Antunes et al., 2011, 2012) |
| CodY | 160<sup>c</sup> | Protein-DNA binding, transcriptomics | (Dineen et al., 2010) |
| Fur | 19 | Transcriptomics, *in silico*<sup>b</sup> | (Berges et al., 2018; Dubois et al., 2016) |
| PrdR | 181 | Transcriptomics | (Bouillaut et al., 2019) |
| SigB | 57 | Transcriptomics | (Kint et al., 2017) |
| SigD | 155 | Transcriptomics | (El Meouche et al., 2013) |
| SigE | 96 | Transcriptomics | (Saujet et al., 2013) |
| SigF | 25 |  |  |
| SigG | 46 |  |  |
| SigH | 40 | Transcriptomics, *in silico*<sup>b</sup> | (Saujet et al., 2011) |
| SigK | 54 | Transcriptomics | (Saujet et al., 2013) |
| SigL | 46<sup>d</sup> | Transcriptomics, *in silico*<sup>b</sup> | (Soutourina et al., 2020) |
| Spo0A | 276<sup>e</sup> | Transcriptomics | (Fimlaid et al., 2013) |

<sup>a</sup>Only genes classified as CcpA-dependent (in the presence or absence of glucose) by Antunes et al. 2012 were included
<sup>b</sup>*In silico* search of the binding motif of the corresponding regulator within the genes and their promoter sequences
<sup>c</sup>Only genes negatively influenced by CodY were included
<sup>d</sup>Only genes with SigL motif and downregulated in *sigL* deletion were included
<sup>e</sup>Only genes positively influenced by Spo0A were included

**Table S3.** General properties of modules associated with the pathogenicity loci

| Gene | Locus tag<sup>a</sup> | Module<sup>b</sup> | Residual | Functional enrichment<sup>c,d</sup> | Enriched TF regulons<sup>c,d</sup> | Inferelator predicted regulators<sup>a,c</sup> |
| --- | --- | --- | --- | --- | --- | --- |
| *tcdR* | CD_06590 | 31 | 0.46 | NF | NF | SigG, SpoVT |
|  |  | 336 | 0.54 | NF | NF | FapR, CD_16930 |
| *tcdB* | CD_06600 | 284 | 0.63 | NF | NF | CD_06930, CD_17820 |
|  |  | 397 | 0.49 | NF | Spo0A, SigE | SpoVT, CD_06290 |
| *tcdE* | CD_06610 | 31 | 0.46 | NF | NF | SigG, SpoVT |
|  |  | 138 | 0.61 | NF | NF | CD_06290, CD_12390, SpoVT |
| *tcdA* | CD_06630 | 182 | 0.56 | ATP synthesis | CodY, PrdR | FapR, CD_20480, SigG |
|  |  | 264 | 0.71 | NF | NF | SigV |
| *tcdC* | CD_06640 | 146 | 0.52 | NF | NF | CD_27320, CD630_05790, PhoU |
|  |  | 395 | 0.51 | NF | NF | CD_18100, CD_35440 |

<sup>a</sup>The 'CD630\_' segment of the locus tag was replace with 'CD\_'
<sup>b</sup>Modules that contain the corresponding member of the pathogenicity loci
<sup>c</sup>Refers to the information shown for the corresponding module on the Cdiff Web Portal
<sup>d</sup>'NF' indicates that no enrichment for functional terms or TF regulons was detected

**Table S4.** Locus tag of uncharacterized genes associated with selected functional terms

| Function Term | Enriched Modules | Enriched TF regulons | Uncharacterized genes |
| --- | --- | --- | --- |
| Other Sugar-family Transporters | 43 | NA | CD630_04371, CD630_09341, CD630_12100, CD630_13011, CD630_13781, CD630_13782, CD630_15970, CD630_17720, CD630_20101, CD630_20751, CD630_29661, CD630_31110 |
| Sporulation | 206, 251 | Spo0A, SigE, SigG | CD630_01310, CD630_02140, CD630_10650, CD630_10660, CD630_19280, CD630_23740, CD630_23750, CD630_24430, CD630_26340, CD630_32590 |
| Diffocin Locus Proteins | 136 | NA | CD630_12420, CD630_12430, CD630_12440, CD630_13760, CD630_14150, CD630_14160 |
| Multi-drug Transport Systems | 29 | NA | CD630_03790, CD630_03800, CD630_10991, CD630_33681 |
| Aminobutanoate degradation | 48 | CodY, PrdR | CD630_05100, CD630_05101, CD630_08760, CD630_08780 |
| Amino-acyl tRNA Synthesis and Modification | 333 | NA | CD630_12860, CD630_26940 |
| Wood-Ljungdahl Pathway | 159, 168 | CcpA, GlnF | CD630_20210, CD630_29280 |
| Xanthine, Hypoxanthine Transport and Metabolism | 145 | NA | CD630_17570, CD630_22580 |
| Flagellar Motility | 296 | SigD | CD630_12920, CD630_13900 |
| Ribosome Synthesis and Modification | 317 | NA | CD630_22121 |
| Cobalamin Biosynthesis | 157 | NA | CD630_04760 |
| Gram+ Peptidoglycan and Teichoic Acid | 121 | NA | CD630_15620 |
| Chemotaxis | 342 | SigD | CD630_32720 |

**Table S5.** Model predicted essential genes in vivo

| Locus tag | Gene name | Gene product/function | Pathway | *In silico* predictions |  | Experiment |
| --- | --- | --- | --- | --- | --- | --- |
|  |  |  |  | Essentiality *in vivo* | Essentiality nutrient-rich broth *in vitro* | Essentiality nutrient-rich broth *in vitro* (Tn-seq) |
| CD630_23430 | cat1 | succinyl-CoA:coenzyme A transferase | Butanoate fermentation; Methionine Biosynthesis; Central carbon metabolism | Essential | Non-Essential | Non-Essential |
| CD630_34400 | G12WB-3619 | glycoside hydrolase-type carbohydrate-binding protein | Glycolysis | Essential | Non-Essential | Non-Essential |
| CD630_12240 | pupG | purine nucleoside phosphorylase | Purine metabolism | Essential | Non-Essential | Non-Essential |
| CD630_07190 | fchA | methenyltetrahydrofolate cyclohydrolase | One carbon pool by folate; Wood-Ljungdahl pathway | Essential | Non-Essential | Non-Essential |
| CD630_12230 | drm | phosphopentomutase | Pentose-phosphate pathway; Nucleotide interconversion | Essential | Non-Essential | Non-Essential |
| CD630_15020 | deoC | deoxyribose-phosphate aldolase | Pentose-phosphate pathway; Nucleotide interconversion | Essential | Non-Essential | Non-Essential |
| CD630_12200 | nudF | NUDIX family hydrolase | Purine metabolism | Essential | Non-Essential | Non-Essential |
| CD630_23300 | xpt | xanthine phosphoribosyltransferase | Purine metabolism | Essential | Non-Essential | Non-Essential |
| CD630_05570 | G12WB-669 | uridine kinase | Pyrimidine metabolism | Essential | Non-Essential | Non-Essential |
| CD630_18160 | cmk | cytidylate kinase | Pyrimidine metabolism | Essential | Non-Essential | Non-Essential |

## Notes

### Summary of Updates

Integration of the transcriptional and metabolic networks into a Phenotype of Regulatory influences Integrated with Metabolism and Environment (PRIME) model to evaluate context-specific TF essentiality.

## REFERENCES

1. Aktories, K. (2011). Bacterial protein toxins that modify host regulatory GTPases. Nat. Rev. Microbiol. 9, 487–498.

2. Antunes, A., Martin-Verstraete, I., and Dupuy, B. (2011). CcpA-mediated repression of Clostridium difficile toxin gene expression. Mol. Microbiol. 79, 882–899.

3. Antunes, A., Camiade, E., Monot, M., Courtois, E., Barbut, F., Sernova, N. V, Rodionov, D.A., Martin-Verstraete, I., and Dupuy, B. (2012). Global transcriptional control by glucose and carbon regulator CcpA in *Clostridium difficile*. Nucleic Acids Res. 40, 10701–10718.

4. Arndt, D., Grant, J.R., Marcu, A., Sajed, T., Pon, A., Liang, Y., and Wishart, D.S. (2016). PHASTER: a better, faster version of the PHAST phage search tool. Nucleic Acids Res. 44, W16--W21.

5. Arrieta-Ortiz, M.L., Hafemeister, C., Bate, A.R., Chu, T., Greenfield, A., Shuster, B., Barry, S.N., Gallitto, M., Liu, B., Kacmarczyk, T., et al. (2015). An experimentally supported model of the *Bacillus subtilis* global transcriptional regulatory network. Mol. Syst. Biol. 11.

6. Arrieta-Ortiz, M.L., Hafemeister, C., Shuster, B., Baliga, N.S., Bonneau, R., and Eichenberger, P. (2020). Inference of Bacterial Small RNA Regulatory Networks and Integration with Transcription Factor-Driven Regulatory Networks. Msystems 5.

7. Bailey, T.L., Johnson, J., Grant, C.E., and Noble, W.S. (2015). The MEME suite. Nucleic Acids Res. 43, W39--W49.

8. Barrett, T., Wilhite, S.E., Ledoux, P., Evangelista, C., Kim, I.F., Tomashevsky, M., Marshall, K.A., Phillippy, K.H., Sherman, P.M., Holko, M., et al. (2012). NCBI GEO: archive for functional genomics data sets—update. Nucleic Acids Res. 41, D991--D995.

9. Becker, S.A., and Palsson, B.O. (2008). Context-specific metabolic networks are consistent with experiments. PLoS Comput Biol 4, e1000082.

10. Berges, M., Michel, A.-M., Lassek, C., Nuss, A.M., Beckstette, M., Dersch, P., Riedel, K., Sievers, S., Becher, D., Otto, A., et al. (2018). Iron regulation in *Clostridioides difficile*. Front. Microbiol. 9, 3183.

11. Bonneau, R., Facciotti, M.T., Reiss, D.J., Schmid, A.K., Pan, M., Kaur, A., Thorsson, V., Shannon, P., Johnson, M.H., Bare, J.C., et al. (2007). A predictive model for transcriptional control of physiology in a free living cell. Cell 131, 1354–1365.

12. Bouillaut, L., Self, W.T., and Sonenshein, A.L. (2013). Proline-dependent regulation of *Clostridium difficile* Stickland metabolism. J. Bacteriol. 195, 844–854.

13. Bouillaut, L., Dubois, T., Francis, M.B., Daou, N., Monot, M., Sorg, J.A., Sonenshein, A.L., and Dupuy, B. (2019). Role of the global regulator Rex in control of NAD+-regeneration in *Clostridioides* (*Clostridium*) *difficile*. Mol. Microbiol. 111, 1671–1688.

14. Bradshaw, W.J., Kirby, J.M., Roberts, A.K., Shone, C.C., and Acharya, K.R. (2017). The molecular structure of the glycoside hydrolase domain of Cwp19 from *Clostridium difficile*. FEBS J. 284, 4343–4357.

15. Bradshaw, W.J., Bruxelle, J.-F., Kovacs-Simon, A., Harmer, N.J., Janoir, C., Péchiné, S., Acharya, K.R., and Michell, S.L. (2019). Molecular features of lipoprotein CD0873: A potential vaccine against the human pathogen *Clostridioides difficile*. J. Biol. Chem. 294, 15850–15861.

16. Brooks, A.N., Turkarslan, S., Beer, K.D., Yin Lo, F., and Baliga, N.S. (2011). Adaptation of cells to new environments. Wiley Interdiscip. Rev. Syst. Biol. Med. 3, 544–561.

17. Brooks, A.N., Reiss, D.J., Allard, A., Wu, W.-J., Salvanha, D.M., Plaisier, C.L., Chandrasekaran, S., Pan, M., Kaur, A., and Baliga, N.S. (2014). A system-level model for the microbial regulatory genome. Mol. Syst. Biol. 10, 740–740.

18. Carter, G.P., Douce, G.R., Govind, R., Howarth, P.M., Mackin, K.E., Spencer, J., Buckley, A.M., Antunes, A., Kotsanas, D., Jenkin, G.A., et al. (2011). The anti-sigma factor TcdC modulates hypervirulence in an epidemic BI/NAP1/027 clinical isolate of *Clostridium difficile*. PLoS Pathog 7, e1002317.

19. Cartman, S.T., Kelly, M.L., Heeg, D., Heap, J.T., and Minton, N.P. (2012). Precise manipulation of the *Clostridium difficile* chromosome reveals a lack of association between the *tcdC* genotype and toxin production. Appl. Environ. Microbiol. 78, 4683–4690.

20. Consortium, T.U. (2017). UniProt: the universal protein knowledgebase. Nucleic Acids Res. 45, D158--D169.

21. Dehal, P.S., Joachimiak, M.P., Price, M.N., Bates, J.T., Baumohl, J.K., Chivian, D., Friedland, G.D., Huang, K.H., Keller, K., Novichkov, P.S., et al. (2010). MicrobesOnline: an integrated portal for comparative and functional genomics. Nucleic Acids Res. 38, D396–400.

22. Dembek, M., Barquist, L., Boinett, C.J., Cain, A.K., Mayho, M., Lawley, T.D., Fairweather, N.F., and Fagan, R.P. (2015). High-throughput analysis of gene essentiality and sporulation in *Clostridium difficile*. MBio 6, e02383--14.

23. Dineen, S.S., Villapakkam, A.C., Nordman, J.T., and Sonenshein, A.L. (2007). Repression of *Clostridium difficile* toxin gene expression by CodY. Mol. Microbiol. 66, 206–219.

24. Dineen, S.S., McBride, S.M., and Sonenshein, A.L. (2010). Integration of metabolism and virulence by *Clostridium difficile* CodY. J. Bacteriol. 192, 5350–5362.

25. Dingle, T.C., Mulvey, G.L., and Armstrong, G.D. (2011). Mutagenic analysis of the *Clostridium difficile* flagellar proteins, FliC and FliD, and their contribution to virulence in hamsters. Infect. Immun. 79, 4061–4067.

26. Dubois, T., Dancer-Thibonnier, M., Monot, M., Hamiot, A., Bouillaut, L., Soutourina, O., Martin-Verstraete, I., and Dupuy, B. (2016). Control of *Clostridium difficile* physiopathology in response to cysteine availability. Infect. Immun. 84, 2389–2405.

27. Edwards, A.N., Tamayo, R., and McBride, S.M. (2016). A novel regulator controls *Clostridium difficile* sporulation, motility and toxin production. Mol. Microbiol. 100, 954–971.

28. Elena, S.F., and Lenski, R.E. (2003). Evolution experiments with microorganisms: the dynamics and genetic bases of adaptation. Nat. Rev. Genet. 4, 457–469.

29. Fimlaid, K.A., Bond, J.P., Schutz, K.C., Putnam, E.E., Leung, J.M., Lawley, T.D., and Shen, A. (2013). Global analysis of the sporulation pathway of *Clostridium difficile*. PLoS Genet. 9.

30. Galperin, M.Y., Makarova, K.S., Wolf, Y.I., and Koonin, E. V (2015). Expanded microbial genome coverage and improved protein family annotation in the COG database. Nucleic Acids Res. 43, D261--D269.

31. Girinathan, B.P., DiBenedetto, N., Worley, J.N., Peltier, J., Arrieta-Ortiz, M., Immanuel, S.R.C., Lavin, Ri., Delaney, M.L., Cummins, C., Onderdonk, A.B., et al. (2021). The mechanisms of in vivo commensal control of Clostridioides difficile virulence. BioRxiv 2020.01.04.894915.

32. Gößner, A.S., Picardal, F., Tanner, R.S., and Drake, H.L. (2008). Carbon metabolism of the moderately acid-tolerant acetogen *Clostridium drakei* isolated from peat. FEMS Microbiol. Lett. 287, 236–242.

33. Govind, R., and Dupuy, B. (2012). Secretion of *Clostridium difficile* toxins A and B requires the holin-like protein TcdE. PLoS Pathog 8, e1002727.

34. Heirendt, L., Arreckx, S., Pfau, T., Mendoza, S.N., Richelle, A., Heinken, A., Haraldsdóttir, H.S., Wachowiak, J., Keating, S.M., Vlasov, V., et al. (2019). Creation and analysis of biochemical constraint-based models using the COBRA Toolbox v. 3.0. Nat. Protoc. 14, 639–702.

35. Immanuel, S.R.C., Arrieta-Ortiz, M.L., Ruiz, R.A., Pan, M., de Lomana, A.L.G., Peterson, E.J.R., and Baliga, N.S. (2021). Quantitative prediction of conditional vulnerabilities in regulatory and metabolic networks of *Mycobacterium tuberculosis*. BioRxiv.

36. Jackson, S., Calos, M., Myers, A., and Self, W.T. (2006). Analysis of proline reduction in the nosocomial pathogen *Clostridium difficile*. J. Bacteriol. 188, 8487–8495.

37. Janoir, C., Denève, C., Bouttier, S., Barbut, F., Hoys, S., Caleechum, L., Chapetón-Montes, D., Pereira, F.C., Henriques, A.O., Collignon, A., et al. (2013). Adaptive strategies and pathogenesis of *Clostridium difficile* from *in vivo* transcriptomics. Infect. Immun. 81, 3757–3769.

38. Janvilisri, T., Scaria, J., and Chang, Y.-F. (2010). Transcriptional profiling of *Clostridium difficile* and Caco-2 cells during infection. J. Infect. Dis. 202, 282–290.

39. Jenior, M.L., Leslie, J.L., Young, V.B., and Schloss, P.D. (2017). *Clostridium difficile* colonizes alternative nutrient niches during infection across distinct murine gut microbiomes. Msystems 2.

40. Kanehisa, M., Furumichi, M., Tanabe, M., Sato, Y., and Morishima, K. (2017). KEGG: new perspectives on genomes, pathways, diseases and drugs. Nucleic Acids Res. 45, D353--D361.

41. Karasawa, T., Ikoma, S., Yamakawa, K., and Nakamura, S. (1995). A defined growth medium for *Clostridium difficile*. Microbiology 141, 371–375.

42. Kashaf, S.S., Angione, C., and Lió, P. (2017). Making life difficult for *Clostridium difficile*: augmenting the pathogen’s metabolic model with transcriptomic and codon usage data for better therapeutic target characterization. BMC Syst. Biol. 11, 25.

43. Kint, N., Janoir, C., Monot, M., Hoys, S., Soutourina, O., Dupuy, B., and Martin-Verstraete, I. (2017). The alternative sigma factor σ^B^ plays a crucial role in adaptive strategies of *Clostridium difficile* during gut infection. Environ. Microbiol. 19, 1933–1958.

44. Larocque, M., Chénard, T., and Najmanovich, R. (2014). A curated *C. difficile* strain 630 metabolic network: prediction of essential targets and inhibitors. BMC Syst. Biol. 8, 117.

45. Love, M.I., Huber, W., and Anders, S. (2014). Moderated estimation of fold change and dispersion for RNA-seq data with DESeq2. Genome Biol. 15, 550.

46. Mani, N., and Dupuy, B. (2001). Regulation of toxin synthesis in *Clostridium difficile* by an alternative RNA polymerase sigma factor. Proc. Natl. Acad. Sci. 98, 5844–5849.

47. Martin-Verstraete, I., Peltier, J., and Dupuy, B. (2016). The regulatory networks that control *Clostridium difficile* toxin synthesis. Toxins (Basel). 8, 153.

48. Matamouros, S., England, P., and Dupuy, B. (2007). *Clostridium difficile* toxin expression is inhibited by the novel regulator TcdC. Mol. Microbiol. 64, 1274–1288.

49. McDonald, J.A.K., Mullish, B.H., Pechlivanis, A., Liu, Z., Brignardello, J., Kao, D., Holmes, E., Li, J. V, Clarke, T.B., Thursz, M.R., et al. (2018). Inhibiting growth of *Clostridioides difficile* by restoring valerate, produced by the intestinal microbiota. Gastroenterology 155, 1495–1507.

50. El Meouche, I., Peltier, J., Monot, M., Soutourina, O., Pestel-Caron, M., Dupuy, B., and Pons, J.-L. (2013). Characterization of the SigD regulon of *C. difficile* and its positive control of toxin production through the regulation of *tcdR*. PLoS One 8.

51. Monegro, A.F., and Regunath, H. (2018). Hospital acquired infections. In StatPearls, (StatPearls Publishing)

52. Monot, M., Boursaux-Eude, C., Thibonnier, M., Vallenet, D., Moszer, I., Medigue, C., Martin-Verstraete, I., and Dupuy, B. (2011). Reannotation of the genome sequence of Clostridium difficile strain 630.

53. Moretto, M., Sonego, P., Dierckxsens, N., Brilli, M., Bianco, L., Ledezma-Tejeida, D., Gama-Castro, S., Galardini, M., Romualdi, C., Laukens, K., et al. (2016). COLOMBOS v3.0: leveraging gene expression compendia for cross-species analyses. Nucleic Acids Res. 44, D620–3.

54. Nawrocki, K.L., Wetzel, D., Jones, J.B., Woods, E.C., and McBride, S.M. (2018). Ethanolamine is a valuable nutrient source that impacts *Clostridium difficile* pathogenesis. Environ. Microbiol. 20, 1419–1435.

55. Neumann-Schaal, M., Metzendorf, N.G., Troitzsch, D., Nuss, A.M., Hofmann, J.D., Beckstette, M., Dersch, P., Otto, A., and Sievers, S. (2018). Tracking gene expression and oxidative damage of O2-stressed *Clostridioides difficile* by a multi-omics approach. Anaerobe 53, 94–107.

56. Nguyen, N.T.T., Contreras-Moreira, B., Castro-Mondragon, J.A., Santana-Garcia, W., Ossio, R., Robles-Espinoza, C.D., Bahin, M., Collombet, S., Vincens, P., Thieffry, D., et al. (2018). RSAT 2018: regulatory sequence analysis tools 20th anniversary. Nucleic Acids Res. 46, W209-W214.

57. Orth, J.D., Thiele, I., and Palsson, B.Ø. (2010). What is flux balance analysis? Nat. Biotechnol. 28, 245–248.

58. Paquette, S.M., Leinonen, K., and Longabaugh, W.J.R. (2016). BioTapestry now provides a web application and improved drawing and layout tools. F1000Research 5.

59. Peltier, J., Hamiot, A., Garneau, J.R., Boudry, P., Maikova, A., Hajnsdorf, E., Fortier, L.-C., Dupuy, B., and Soutourina, O. (2020). Type I toxin-antitoxin systems contribute to the maintenance of mobile genetic elements in *Clostridioides difficile*. Commun. Biol. 3, 1–13.

60. Peterson, E.J.R., Reiss, D.J., Turkarslan, S., Minch, K.J., Rustad, T., Plaisier, C.L., Longabaugh, W.J.R., Sherman, D.R., and Baliga, N.S. (2014). A high-resolution network model for global gene regulation in *Mycobacterium tuberculosis*. Nucleic Acids Res. 42, 11291–11303.

61. Reiss, D.J., Plaisier, C.L., Wu, W.-J., and Baliga, N.S. (2015). cMonkey2: Automated, systematic, integrated detection of co-regulated gene modules for any organism. Nucleic Acids Res. 43, e87.

62. Riedel, T., Bunk, B., Thürmer, A., Spröer, C., Brzuszkiewicz, E., Abt, B., Gronow, S., Liesegang, H., Daniel, R., and Overmann, J. (2015). Genome resequencing of the virulent and multidrug-resistant reference strain *Clostridium difficile* 630. Genome Announc. 3, e00276--15.

63. Saujet, L., Monot, M., Dupuy, B., Soutourina, O., and Martin-Verstraete, I. (2011). The key sigma factor of transition phase, SigH, controls sporulation, metabolism, and virulence factor expression in *Clostridium difficile*. J. Bacteriol. 193, 3186–3196.

64. Saujet, L., Pereira, F.C., Serrano, M., Soutourina, O., Monot, M., Shelyakin, P. V, Gelfand, M.S., Dupuy, B., Henriques, A.O., and Martin-Verstraete, I. (2013). Genome-wide analysis of cell type-specific gene transcription during spore formation in *Clostridium difficile*. PLoS Genet 9, e1003756.

65. Seemann, T. (2014). Prokka: rapid prokaryotic genome annotation. Bioinformatics 30, 2068– 2069.

66. Soutourina, O., Dubois, T., Monot, M., Shelyakin, P. V, Saujet, L., Boudry, P., Gelfand, M.S., Dupuy, B., and Martin-Verstraete, I. (2020). Genome-Wide Transcription Start Site Mapping and Promoter Assignments to a Sigma Factor in the Human Enteropathogen *Clostridioides difficile*. Front. Microbiol. 11, 1939.

67. Soutourina, O.A., Monot, M., Boudry, P., Saujet, L., Pichon, C., Sismeiro, O., Semenova, E., Severinov, K., Le Bouguenec, C., Coppée, J.-Y., et al. (2013). Genome-wide identification of regulatory RNAs in the human pathogen *Clostridium difficile*. PLoS Genet 9, e1003493.

68. Steglich, M., Hofmann, J.D., Helmecke, J., Sikorski, J., Spröer, C., Riedel, T., Bunk, B., Overmann, J., Neumann-Schaal, M., and Nübel, U. (2018). Convergent loss of ABC transporter genes from *Clostridioides difficile* genomes is associated with impaired tyrosine uptake and p-cresol production. Front. Microbiol. 9, 901.

69. Szklarczyk, D., Morris, J.H., Cook, H., Kuhn, M., Wyder, S., Simonovic, M., Santos, A., Doncheva, N.T., Roth, A., Bork, P., et al. (2016). The STRING database in 2017: quality-controlled protein--protein association networks, made broadly accessible. Nucleic Acids Res. gkw937.

70. Tatusova, T., DiCuccio, M., Badretdin, A., Chetvernin, V., Nawrocki, E.P., Zaslavsky, L., Lomsadze, A., Pruitt, K.D., Borodovsky, M., and Ostell, J. (2016). NCBI prokaryotic genome annotation pipeline. Nucleic Acids Res. 44, 6614–6624.

71. Theriot, C.M., Koenigsknecht, M.J., Carlson Jr, P.E., Hatton, G.E., Nelson, A.M., Li, B., Huffnagle, G.B., Li, J.Z., and Young, V.B. (2014). Antibiotic-induced shifts in the mouse gut microbiome and metabolome increase susceptibility to *Clostridium difficile* infection. Nat. Commun. 5, 3114.

72. Underwood, S., Guan, S., Vijayasubhash, V., Baines, S.D., Graham, L., Lewis, R.J., Wilcox, M.H., and Stephenson, K. (2009). Characterization of the sporulation initiation pathway of *Clostridium difficile* and its role in toxin production. J. Bacteriol. 191, 7296–7305.

73. Vallenet, D., Calteau, A., Cruveiller, S., Gachet, M., Lajus, A., Josso, A., Mercier, J., Renaux, A., Rollin, J., Rouy, Z., et al. (2017). MicroScope in 2017: an expanding and evolving integrated resource for community expertise of microbial genomes. Nucleic Acids Res. 45, D517--D528.

74. Vemuri, R.C., Gundamaraju, R., Shinde, T., and Eri, R. (2017). Therapeutic interventions for gut dysbiosis and related disorders in the elderly: antibiotics, probiotics or faecal microbiota transplantation? Benef. Microbes 8, 179–192.

75. Walter, B.M., Rupnik, M., Hodnik, V., Anderluh, G., Dupuy, B., Paulič, N., Žgur-Bertok, D., and Butala, M. (2014). The LexA regulated genes of the *Clostridium difficile*. BMC Microbiol. 14, 88.

76. Wattam, A.R., Abraham, D., Dalay, O., Disz, T.L., Driscoll, T., Gabbard, J.L., Gillespie, J.J., Gough, R., Hix, D., Kenyon, R., et al. (2014). PATRIC, the bacterial bioinformatics database and analysis resource. Nucleic Acids Res. 42, D581--D591.

77. Wolfe, C.J., Kohane, I.S., and Butte, A.J. (2005). Systematic survey reveals general applicability of “guilt-by-association” within gene coexpression networks. BMC Bioinformatics 6, 227.

78. Woods, E.C., Nawrocki, K.L., Suárez, J.M., and McBride, S.M. (2016). The *Clostridium difficile* Dlt pathway is controlled by the extracytoplasmic function sigma factor σ^V^ in response to lysozyme. Infect. Immun. 84, 1902–1916.

